# Histopathologic Evaluation and Single-Cell Spatial Transcriptomics of the Colon Reveal Cellular and Molecular Abnormalities Linked to J-Pouch Failure in Patients with Inflammatory Bowel Disease

**DOI:** 10.1101/2025.01.27.635092

**Authors:** Andrea D. Olivas, Paul Chak Mou Ngai, Emily Schahrer, Kinga S. Olortegui, John F. Cursio, Shintaro Akiyama, Eugene B. Chang, Le Shen, Konstantin Umanskiy, David T. Rubin, David Zemmour, Christopher R. Weber

## Abstract

**Background and Aims:** Total abdominal colectomy (TAC) with a staged ileal pouch-anal anastomosis (IPAA) is a common surgical treatment for ulcerative colitis (UC). However, a significant percentage of patients experience pouch failure, leading to morbidity. This retrospective case-control study identified histopathological features of the TAC specimen associated with pouch failure and investigated the molecular mechanisms of this susceptibility using single-cell spatial transcriptomics.

**Methods:** We analyzed a cohort of 417 patients who underwent IPAA between 2000-2010 at the University of Chicago Medical Center for up to 18 years. Histological examination of TAC specimens focused on disease activity, depth of inflammation, and specific features, including granulomas and deep ulcers. A subset of patients was profiled using single-cell spatial transcriptomics to map gene expression and immune cell interactions in relation to the risk of pouch failure.

**Results:** The 18-year pouch failure risk was 23%, with post-procedure diagnosis of CD as a major risk factor (HR = 4.3, 95% CI: 2.3–8.1) as well as high-risk histologic features, including deep chronic inflammation (HR = 21, 95% CI: 11-41) and severe disease activity (HR = 14, 95% CI: 5.7-32) in TAC specimens. Spatial transcriptomics showed immune infiltration of T and myeloid cells, reduced myocyte-glial interactions, and cytokine signaling pathways such as IL-10, IL-1β, and type I/II interferons, associated with an increased risk of pouch failure.

**Conclusion:** Histological features and spatial molecular profiling are predictive of IPAA failure. These findings support the use of histologic evaluation and targeted molecular analysis of the TAC specimen to identify high-risk patients and improve IPAA outcomes.

## Introduction

Surgical treatment of ulcerative colitis (UC) involves colon removal, with restorative proctectomy and J-pouch formation reestablishing intestinal continuity.^1, 2^ When first introduced, proctocolectomy with IPAA was performed either as a single operation or a two-stage procedure employing temporary proximal diversion with loop ileostomy followed by interval ileostomy reversal.^3, 4^ Over the last several decades, as the medical management of UC became the mainstay of treatment, patients in whom medical therapy failed and who presented for surgery were noted to have more severely active and resistant disease. Accordingly, surgical techniques evolved to accommodate this change in patient population. As data regarding postoperative complications and patient outcomes have become available, there has been a steady trend favoring a three-stage procedure involving total abdominal colectomy (TAC) with end ileostomy as the first stage, followed by interval restorative proctectomy and IPAA with diverting loop ileostomy (second stage), and subsequent ileostomy reversal as the final stage.

Despite improved techniques, IPAA remains linked to morbidity and pouch failure. Nearly half of patients with a pouch experience at least one episode of pouchitis, and up to 12% will require a pouch excision or permanent diversion within 10 years due to pouch failure ^5–7^. Crohn’s-like features, such as ileal stenosis and perianal complications, are a common cause of pouch failure.

Identifying high-risk IPAA patients is essential^8^ and several studies have identified risk factors for pouch failure. The preoperative diagnosis of Crohn’s disease (CD) is a relative contraindication to IPAA, considering the overall risk of pouch failure approaching 50% compared to 10% in ulcerative colitis (UC) patients ^9–16^. The Chicago Classification of Pouch Endoscopy^17^ has defined seven distinct phenotypes with different pouch failure rates, and other studies have incorporated abnormal anal manometry, pelvic sepsis, and anastomotic issues for risk stratification. However, these predictive models incorporate post-IPAA factors, requiring patients to undergo pouch formation before a comprehensive risk assessment can be conducted^18,19^. A significant benefit of a staged IPAA procedure is the opportunity for careful histologic analysis of the initial colectomy specimen, prior to interval pouch formation.

Therefore, this study aimed to identify histologic features in the TAC specimen that correlate with subsequent pouch failure. We found that approximately 10% of IPAA patients in our cohort experienced pouch failure, with many later reclassified as CD, and that failure was significantly associated with high-risk histologic features in TAC specimens, including disease severity, deep chronic inflammation, and non-UC histologic features.

The second aim of the study was to identify biological mechanisms of pouch failure susceptibility. Previous studies have demonstrated that the interaction between commensal bacteria and the mucosal immune system plays a significant role in the pathogenesis of pouchitis^20^. In particular, gut dysbiosis associated with pouchitis has been shown to drive heightened T-cell proliferation and activation^21, 22^. Shifts in macrophage and T cell subpopulations have been observed with altered cytokine profiles, such as an increase in pro-inflammatory cytokines like TNF-α, IL-1β, and IL-6, and a reduction in immunoregulatory cytokines such as IL-2, and IL-10^23, 24^. However, most of these studies have been limited to analyses of human blood or *in vitro* assays, lacking direct *in situ* examination of immune responses in the tissue and failing to provide predictive insights from TAC tissue prior to pouch formation. We hypothesize that ileal pouch failure in IBD arises from preexisting aberrant cellular interactions present in the TAC specimen. To pinpoint the cellular and immune features predictive of pouch failure, we leveraged recent technical advances in spatial transcriptomics that enable single-molecule RNA detection and single-cell resolution of gene expression across entire tissue sections (CosMx^25^). We show that pouch failure is associated with transmural infiltration and increased interactions of monocytes, macrophages, and T cells, with loss of interactions between glial cells and myocytes. These non-immune cells are key players in maintaining normal motility and structure of the muscularis propria. We further identify layer-specific cellular cross-talks mediated by cytokines and predisposing to pouch failure: type I interferon (IFN) in epithelial cells, IL-1β in the mucosa, and IL-10 in the muscularis propria.

## Materials and methods

### Study Cohort

Patients with IBD who underwent IPAA from 2000-2010 at the University of Chicago Medical Center were identified from a prospectively maintained, IRB-approved surgical database (IRB 16014A). IPAA outcomes were classified as failure (requiring excision or permanent ileostomy) or non-failure, with records reviewed for pouch duration and failure type. Patients with early technical failure (<3 months) or cancer-related complications were excluded (n=3).

### Definition of Crohn’s Disease (CD)

CD was defined to include (1) pre-TAC diagnosis of CD, (2) histologic diagnosis of CD in the TAC specimen, and (3) evolution of ulcerative/indeterminate colitis to a CD phenotype after TAC or IPAA. This definition encompassed “pouchitis with CD-like features” and CD-like strictures of the pre-pouch ileum. Patients were not subcategorized by timing of diagnosis.

### Pouch Survival Analysis

Kaplan-Meier curves and log-rank tests assessed pouch survival (n=414) for CD *vs*. non-CD diagnoses. Cox proportional hazards calculated hazard ratios (HR) and 95% confidence intervals (CI) to quantify pouch failure risk. Inverse probability weighting (IPW) Kaplan-Meier analysis was used to adjust for sampling bias, incorporating severity and depth of inflammation^26^.

### Histologic Evaluation

TAC specimens from 39 IPAA failure patients (out of 41) were histologically analyzed and matched 1:1 with non-failure patients. Matching criteria included age, sex, and number of operative stages. Clinical data included disease duration, biologic use, preoperative *C. difficile* infection, preoperative biopsy diagnosis, and follow-up time. H&E-stained slides of TAC specimens (proximal/distal margins, appendix, and section every 10 cm) were reviewed blindly by two pathologists. Disease severity was scored using published criteria^27^, and the depth of inflammation was graded from 0-4 (0 = normal to 4 = transmural). Non-UC features recorded included epithelioid granulomas (not linked to crypt rupture), ileal inflammation, deep “knife-like” ulcers, right-sided disease, and patchy colonic disease. Histologic findings were correlated with clinical outcomes.

### Single-cell Resolution Spatial Transcriptomics

Sample Processing. Spatial transcriptomics was conducted on 8 FFPE colon samples (4 failure, 4 non-failure). Samples were processed on the CosMx instrument using a 1K Human Universal Panel (974 genes) and a 20-gene custom panel.

Data processing and analysis. Cells were segmented using CellPose^28^. Cells with >5% negative probe counts, <10 transcripts were excluded. 97% of cells were retained for analysis. All samples were integrated with SCVI v1.0.0^29^ and projected in two dimensions using Uniform Manifold Approximation and Projection. Tissue layers were detected with Vesalius^30^. Cell types were clustered and annotated by identifying marker genes for each cluster using FindAllMarkers() (Seurat v5.1.0^31^). Differential gene expression between failure and non-failure samples was analyzed using pseudobulk limma-trend (limma v3.58.1^32^, edgeR v4.0.16^33^). For neighborhood analysis, cell distances were measured using the euclidian distance. K-nearest neighbor analysis was used to quantify cell pairs in tissue samples with EdgeR analysis for statistical comparisons. Cell densities were calculated using alphahull library (V2.5)^34^. Pathway Enrichment Analysis was performed with the pathfindR package (V2.3.1) and the Reactome^35^ datasets. CellChat v2.0^36^ and the Immune Dictionary App^37^ were used to investigate intercellular communication.

### Statistical Analysis

Analysis was conducted using R-3.6.2. Univariate statistical analysis was performed using the Fisher Exact Test or 2-tailed Students *t*-test where appropriate. Additional statistical tests, including logistic regression, are described in Supplementary Methods. *P-value*s or adjusted *P-values* using the Benjamini-Hochberg procedure of less than 5% were deemed significant.

### Detailed experimental methods are available in Supplementary Methods

## Results

### Pouch failure risk reached 23% over 18 years, with CD/CD-like features being a significant risk factor

417 patients with IBD underwent IPAA at the University of Chicago Medical Center from 2000-2010 and were followed for up to 18 years (median 4.6 years) (**Figure 1A**). Three patients were excluded due to technical failure or complications from metastatic lung cancer. Among the remaining patients, 41/414 (9.9%) experienced IPAA failure requiring pouch excision or permanent diverting ileostomy. Accounting for censored events, Kaplan-Meier pouch survival analysis estimated an 18-year pouch failure risk of 23% (95% CI: 14%-30%) (**Figure 1B**). Most IPAA failures were due to CD-like features of the pouch (56%), followed by pouchitis (27%) and poor function (17%).

**Figure 1:**
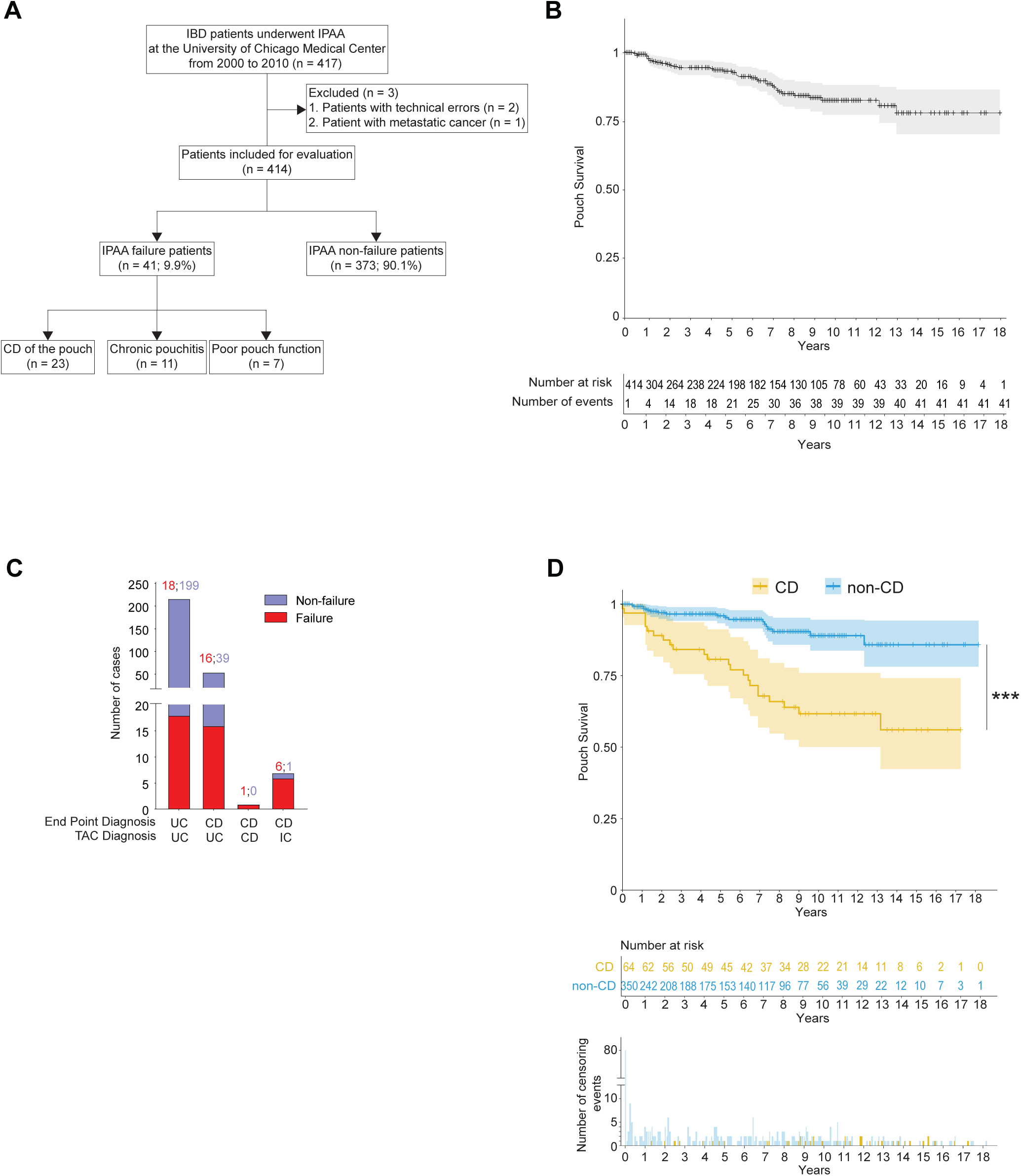
IPAA failure in IBD patients is associated with Crohn’s disease/Crohn’s-like features of the pouch. **(A)** Flowchart with inclusion/exclusion criteria to the retrospective cohort study on IPAA failure in IBD patients treated at The University of Chicago (2000-2010). IPAA outcomes and pouch failure diagnoses presented. **(B)** Kaplan-Meier pouch survival curve (with 95% CI). Tick marks indicate censored data. The table shows the number of patients at risk and cumulative events at each timepoint. **(C)** Bar graphs showing pouch failure proportions by initial TAC diagnosis and subsequent diagnosis revisions for CD, UC, and IC. **(D)** Kaplan-Meier pouch survival curves (with 95% CI) for patients with CD or non-CD of the pouch. Tick marks indicate censored data. The table and barplot show the number of patients at risk and censored events in each group over time. *** *P* < 0.001; log-rank test.

Of the 373 non-failure patients, 239 (64%) had over 24 months of follow-up. Combined with 41 failure cases, 280 patients were analyzed for the correlation between pouch failure and CD diagnosis, assessed post-TAC or during follow-up **(Figure 1C)**. Of these 280 patients, only 1 patient had TAC-consistent Crohn’s and later experienced pouch failure. Patients reclassified from UC or indeterminate colitis (IC) to CD had notably higher pouch failure rates (29% and 86%, respectively) compared to those who remained classified as UC (29% and 8%).

The 18-year cohort enabled time-to-event analysis **(Figure 1D)**. Patients with Crohn’s disease (CD) had a significantly higher pouch failure rate of 44% (95% CI: 26%-58%) compared to 14% (95% CI: 6%-22%) in non-CD patients (HR = 4.3, 95% CI: 2.3-8.1, P = 0.0001, log-rank test). The risk was greatest within the first 10 years, with CD patients showing a 38% pouch survival rate by year 9. Beyond 10 years, the additional risk of pouch failure in CD patients was 6%, similar to non-CD patients. In conclusion, CD of the pouch significantly increases IPAA failure risk, highlighting the need for preoperative identification of CD histologic predictors.

### High-risk histological features of the TAC specimen are associated with pouch failure

To identify histological features predictive of pouch failure, we analyzed TAC specimens from pouch failure cases with available slides (n=39) and a matched cohort of non-failure controls (see Methods and **Table S1**). H&E-stained histologic sections of each colectomy specimen were evaluated by blinded pathologists for disease activity and depth of inflammation every 10 cm, providing a comprehensive map of disease distribution. Disease activity was scored using established criteria^27^: mild (neutrophils in the epithelium), moderate (crypt abscesses), severe (neutrophils with epithelial erosion or ulceration), and quiescent (chronic features without active inflammation) **(Figure 2A, Table S2).** IPAA failures had more areas of severe disease (**Figure 2B)**, with larger ulceration regions than non-failures (24% vs. 11%, P= 0.0036) (**Figure 2C**).

**Figure 2:**
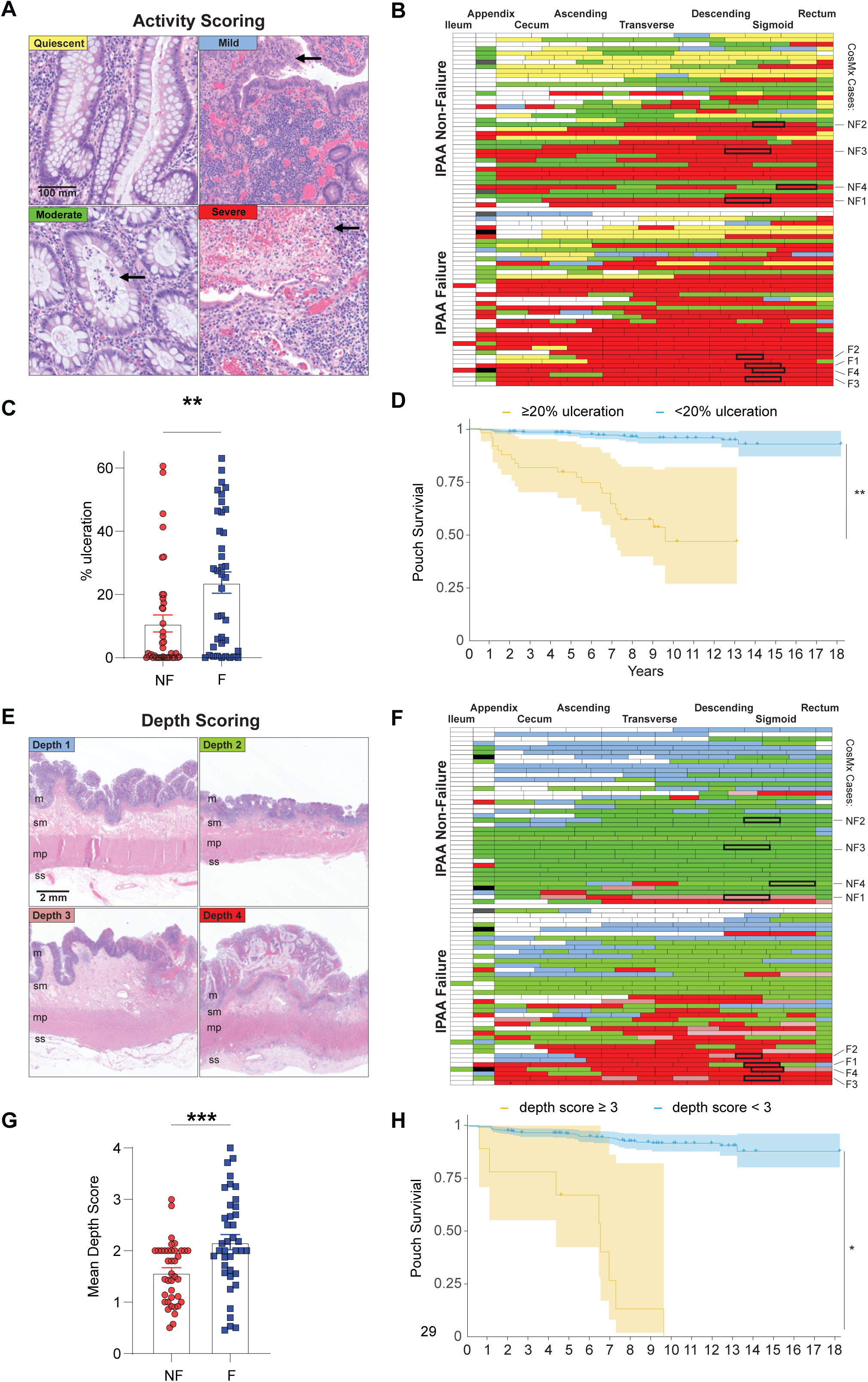
Systematic histologic evaluation of the colectomy specimen (before pouch formation) reveals high-risk histological features associated with pouch failure. **(A)** Representative H&E images of TAC activity scores (normal, quiescent, mild, moderate, severe) with arrows highlighting key features: epithelial neutrophils (mild), crypt abscesses (moderate), and ulceration (severe). **(B)** Heatmap of disease activity every 10 cm along TAC specimens from ileum to rectum, including the appendix when available. Color key: white (normal), yellow (quiescent), blue (mild), green (moderate), red (severe) as shown in A. **(C)** Average ulceration percentage per TAC in IPAA failure vs. non-failure patients. ** *P* < 0.01; two-tailed unpaired Student t-test. **(D)** Kaplan-Meier survival pouch survival curves (with 95% CI) for patients with >20% vs. <20% TAC ulceration. Tick marks indicate censored data. ** *P* < 0.01; log-rank test. **(E)** Representative H&E images showing the depth of chronic inflammation in TAC specimens: score 1 (mucosa, m), score 2 (submucosa, sm), score 3 (muscularis propria, mp), and score 4 (subserosa, ss). **(F)** Heatmap of inflammation depth every 10 cm along TAC specimens from ileum to rectum, including the appendix when available. Color key: white (score 0, normal), blue (score 1), green (score 2), pink (score 3), red (score 4) as shown in E. **(G)** Barplot of average inflammation depth scores in IPAA failure vs. non-failure patients.*** *P* < 0.001; two-tailed unpaired Student t-test. **(H)** Kaplan-Meier survival pouch survival curves (with 95% CI) for patients with TAC depth scores ≥3 vs. <3 (95% CI). Tick marks indicate censored data.** *P* < 0.01; log-rank test.

Inflammation depth was scored from 0 to 4, ranging from no inflammation (0) to inflammation of the mucosa (1), submucosa (2), muscularis propria (3), and pericolic fibroadipose tissue (4) **(Figure 2E, Table S3).** Deeper chronic inflammation was observed in IPAA failures vs. non-failures (average depth of 2.2 vs. 1.6; P=0.0003) (**Figure 2F, G**) and correlated with disease severity (r=0.82, P<10⁻⁹).

To estimate the risk of pouch failure from the TAC, we performed a Kaplan-Meier analysis. Pouch survival was significantly worse when colon ulceration was ≥20% (P=0.001, log-rank test), with a Cox proportional hazard ratio of 14 (95% CI: 5.7-32) and median survival of 9.5 years (**Figure 2D)**. When the mean depth score was ≥3, pouch survival was also significantly affected (IPW Kaplan-Meier, P=0.01, log-rank test), with an HR of 21 (95% CI: 11-41) and median pouch survival of 6.5 years (**Figure 2H)**.

Active disease in the appendix did not differ between groups (59% non-failures vs. 69% failures; P=0.47), nor did deep chronic inflammatory infiltrates (score ≥3) (5.4% non-failures vs. 14% failures; P=0.26, Fisher’s exact test).

In addition to disease severity and inflammation depth, “non-UC” histologic features were recorded (**Table S4**). Deep “knife-like” ulcers were significantly more common in IPAA failures (38% vs. 3% in non-failures; P=0.0001), with 67% (10/15) of these failures linked to CD complications. Transmural inflammation (depth score ≥3) in at least one TAC section was also enriched in IPAA failures at 59% (23/39) vs. 18% (7/39) in non-failures (P=0.0004, Fisher’s exact test), with 57% (13/23) of these failures attributed to CD of the pouch. Other features were noted in failures but too few to reach significance by themselves: epithelioid granulomas (unrelated to crypt rupture) (1/39), active chronic ileal inflammation (3/39), patchy, non-continuous colonic disease (2/39). All six patients later developed CD of the pouch. Right-sided disease with rectal sparing was also found in one IPAA failure (1/39); this patient later experienced IPAA failure due to chronic pouchitis. Combined, at least one high-risk histologic feature was present in 69% (27/39) of IPAA failure TAC specimens vs. 21% (8/39) of non-failures (P<0.0001, Fisher’s exact test).

Highlighting the importance of histological features, stepwise logistic regression assessing pre-IPAA factors (disease duration, preoperative biopsy diagnosis, preoperative biologic use, prior C. difficile colitis treatment, and TAC indication) and TAC histologic characteristics (presence of ≥1 high-risk feature and average ulceration degree) identified having at least one high-risk histologic feature in the TAC specimen as the only significant predictor of pouch failure (OR=9.24, P=0.0024), with the model accurately predicting 79% of cases (c-statistic).

Among the patients for whom we performed histological examination of TAC specimens, a subset underwent pouch endoscopic classification according to the seven phenotypes of the Chicago Classification^17^. This included 13 patients from the failure cohort and 22 from the non-failure cohort. Logistic regression was used to identify associations between histologic features and subsequent pouch endoscopy phenotypes. TAC inflammation depth positively correlated with diffuse pouch inflammation (OR=2.8, 95% CI=1.2–8.4) and pouch-related fistulas (OR=3.3, 95% CI=1.1–13.3). These findings suggest that severe histologic features not only predict pouch failure risk but may also drive the development of distinct pouch phenotypes. These findings underscore the importance of detailed histologic assessment to identify patients at high risk of IPAA failure for individualized care.

### Single-cell spatial transcriptomics reveals deep infiltration of macrophages and T cells in colon tissue, which is associated with a high risk of pouch failure

After evaluating the clinical and histological features predictive of pouch failure, we sought to identify the molecular pathways contributing to it. Using CosMx^25^ a spatial transcriptomic platform that quantifies at the single-cell resolution the expression of 974 genes and protein markers (CD45, pan-cytokeratin (panCK), CD3, and β2m), we profiled TAC specimens from four pouch failure patients and four non-failure patients (CosMx cases F1-F4, NF1-NF4, respectively, highlighted in **Figure 2B, E**). The samples were matched for activity, diagnosis of TAC specimen (all UC), tissue location (descending colon), and profiled on a single slide to limit batch effect. All layers of the colon were represented except for sample F2, in which the mucosa and submucosa were lost during tissue block preparation, and sample F4, which lacked the mucosa due to the severity of the disease **(Figure 3A)**.

**Figure 3:**
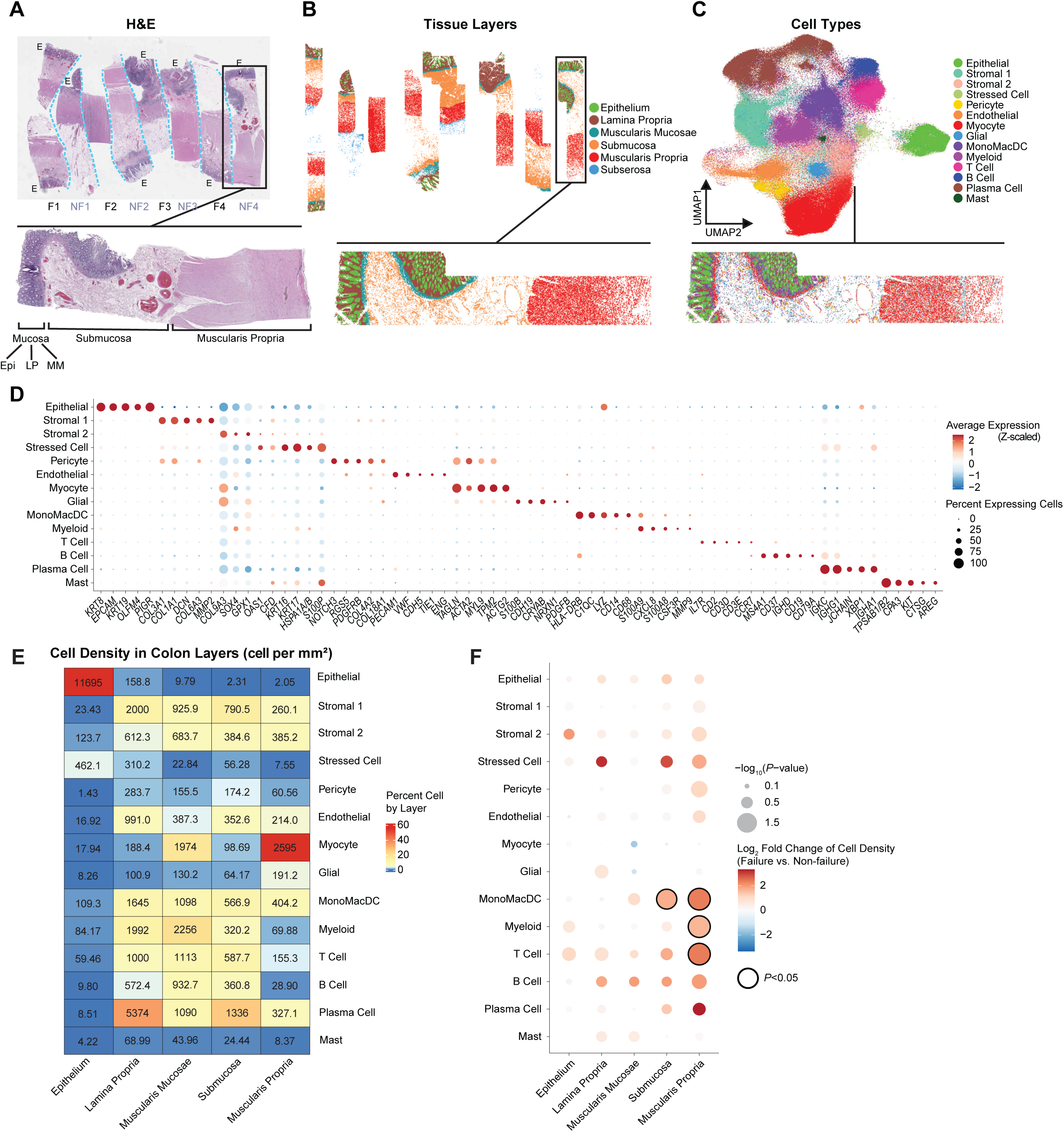
Single-cell spatial transcriptomics reveals extensive macrophage and T-cell infiltration linked to an increased risk of future pouch failure. Four histologically matched cross-sections from failure (F1-4) and non-failure (NF1-4) cases (see Figure 2B, F) were profiled using CosMx. (**A**) H&E-stained adjacent section, with the epithelium marked (E) in each fragment. Inset: NF4 as a representative image. Epi: epithelium, LP: lamina propria, MM: muscularis mucosa (**B**) Tissue layers (in colors) identified using Vesalius (see Methods). Inset: NF4 as a representative image. (**C**) 2D UMAP cell embedding based on gene expression, highlighting 14 major cell types across all samples (in color). The tissue distribution of cell types is shown below, using NF4 as a representative image. (**D**) Dot plot displaying the average expression (in color, scaled) and percentage of cells expressing top marker genes for each cell type. (**E**) Heatmap showing cell type density across colon layers (cells per mm^2^), with color indicating the percentage of each cell type within each layer. (**F**) Dot plot illustrating the fold change in cell density per cell type across layers. Color represents the log2 fold change, and point size is proportional to the *P*-*value*. Statistically significant enrichments (*P* < 0.05, one-sided Wilcoxon rank-sum test) are outlined in black.

After quality control analysis, 352,531 cells were profiled, with an average of 44,066 cells per patient sample (**Table S5**). We used cell membrane protein markers (CD45, pan-cytokeratin, CD3, and β2m) to perform accurate cell segmentation (**Figure S1A**) using CellPose^31^ (see Methods). All 974 genes of the panel were detected, with an average of 193 transcripts and 90 genes per single cell (**Table S5**). We then used SCVI^29^ to integrate the gene expression data across samples (**Figure S1B),** and, using Vesalius^30^ (see Methods), we identified the six different histological components of the colon, which matched H&E sections (**Figure 3A, B**). As expected, the detected epithelium expressed *EPCAM* and panCK and separated from the lamina propria by the expression of *αSMA*, which also highlighted the muscularis propria (**Figure S2A**). The detected submucosa and subserosa were located below the mucosa and muscularis propria, respectively. We then used unsupervised clustering to identify the major cell types of the colon (**Figure 3C**) (see Methods). We annotated them using a multimodal approach, combining their RNA expression, protein expression, and tissue distribution. Epithelial cells mapped spatially to the epithelium (**Figure 3C, E**, density of 11,695 cells/mm^2^) with expression of canonical epithelial marker genes (e.g. *EPCAM*, *KRT9*) (**Figure 3D**, **Figure S2C**). These epithelial cells were also positive for panCK and negative for CD45 (**Figure S2B)**. T cells were most concentrated in the lamina propria (density of 1,000 cells/mm^2^) with CD3 RNA and protein expression (**Figure 3D, E**; **Figure S2B, C**). Other immune cells, such as monocytes, macrophages, and dendritic cells (later referred to as “MonoMacDC”), myeloid cells (granulocytes), B cells, and plasma cells, expressed their expected marker genes, and were positive for CD45 and negative for CD3 (**Figure 3D, E**; **Figure S2B, C**).

Compared to a single-cell RNAseq study without spatial information^38^, spatial transcriptomics had distinct advantages. First, it enabled the profiling of cell populations that are challenging to isolate enzymatically, such as myeloid cells, MonoMacDC cells, endothelial cells, stromal cells, glial cells, and myocytes. Comparing the percentage of cells in single-cell RNAseq and our single-cell spatial transcriptomics dataset, single-cell RNAseq oversampled many immune cells and undersampled parenchymal cells (**Figure S2D)**. For example, stromal cells represented 16% of our data set and less than 4% in a reference single-cell RNAseq dataset of UC,^38^ while T cells represented 6% of cells vs 29% (**Figure S2D**). Secondly, single-cell spatial transcriptomics reports absolute and accurate quantification of cell types in the different colon layers because of the preservation of tissue architecture (**Figure 3E**), which allowed the assessment of differences in cell compositions **(Figure 3F, Figure S3)** and their interactions (see below). Comparing samples from the failure and non-failure cohorts, no significant difference in immune and non-immune cell infiltration was found in the mucosa. Differences were observed in deeper layers, with a peak infiltration of MonoMacDC cells in the muscularis propria (3.7×10^3^ cells/mm^2^ in pouch failures, fold change (FC) 5.10, *P*=0.014, **Figure 3F, Figure S3A)** and T cells (5.1×10^3^ cells/mm^2^ in pouch failures, FC 5.15, *P*=0.014, **Figure 3F, Figure S3B**). The density of other cell types was not significantly different, except for a slightly higher density of granulocytes in the muscularis propria **(Figure S3C-N).** In conclusion, single-cell resolution spatial transcriptomic analysis of pouch failure and non-failure samples revealed deep infiltration of T cells and MonoMacDCs in the tissue of our pouch failure cohort, particularly in the muscularis propria.

### Neighborhood analysis shows increased cellular interactions of immune cells and decreased glial cell-myocyte interactions in samples associated with pouch failure

To gain a deeper understanding of the effects of deep infiltration of T cells and MonoMacDCs in the muscularis propria, we analyzed their spatial distribution, cellular neighbors and created an interaction network among all cells.

Immune infiltration can be diffuse or more organized in granuloma-like aggregates or tertiary lymphoid structures. Cell density heatmaps show that T cell and MonoMacDCs infiltrates was generally diffuse in the submucosa (**Figure 4A**) and muscularis propria (**Figure 4B**). Density distributions of the MonoMacDC population in these tissues were generally unimodal (**Figure 4C, D**), reflecting the diffuse nature of the infiltrate. In addition, the MonoMacDC population was 5.1 times denser in pouch failure relative to non-failure samples (*P*=0.014) (**Figure S3A**). To test whether the infiltrate was organized microscopically, we performed a neighborhood analysis. We quantified the average minimum distance between MonoMacDCs and their neighboring cells, finding that T cells, myeloid cells, and pericytes were closer to MonoMacDCs in pouch failure samples (**Figure 4E**). The average minimum distance of T cells to MonoMacDCs was 52 µm in pouch failures compared to 103 µm in non-failures (*P <* 0.05) (**Figure 4D**). In pouch failure samples, 11% of MonoMacDC were within 10 µm and 92% within 100 µm of T cells, compared to only 2% and 59% in non-failure samples, respectively, suggesting both increased direct (within 10 µm) and indirect (within 100 µm, e.g., cytokine-mediated) cell contact in pouch failure.

**Figure 4.**
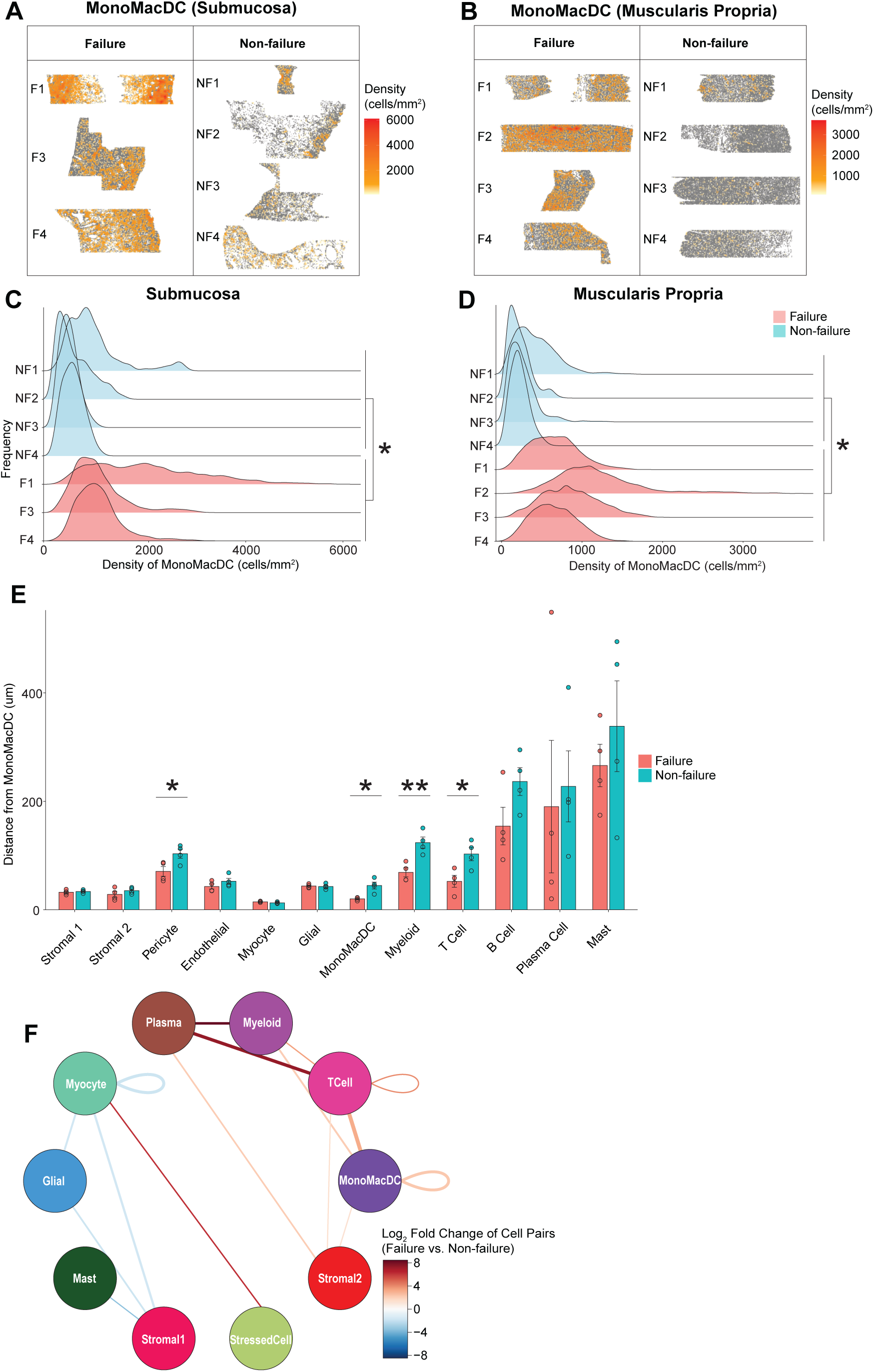
Neighborhood analysis reveals increased infiltration of macrophages and T cells deep into the muscularis propria and a reduction in myocyte-glial cell interactions linked to an elevated risk of future pouch failure. **(A-B)** Heatmaps showing MonoMacDC cell density in the submucosa **(A)** and muscularis propria **(B)** for each sample. Only the muscularis propria was profiled for F3. **(C-D)** Distribution of MonoMacDC density for each patient in the submucosa (C) and muscularis propria (D). * *P <* 0.05, one-sided Wilcoxon rank-sum test. **(E)** Average distance (with SEM) of each MonoMacDC cell to the nearest cell of each major cell type in the muscularis propria. * *P <* 0.05, ** *P <* 0.01, two-tailed t-test. **(F)** Circle plot of enriched cell pairs in pouch failure vs. non-failure, with node size reflecting cell types and edge width the log2 fold change. Only statistically significant edges (*P* < 0.05, moderated two-tailed t-test) are shown.

Having assessed immune interactions in both non-failure and failure tissues, we expanded the neighborhood analysis to understand interactions of all cell types in the muscularis propria and quantified the cell types of all pairs of closest cells in the tissue. Consistent with the immune cell-centric analysis above, there was a striking increase in immune cell interactions in tissues from patients with pouch failure. T cells, MonoMacDCs, plasma cells, and myeloid cells formed a subnetwork of pairwise interactions enriched in these samples (**Figure 4F**). Myocyte interactions with stressed stromal cells were also increased. In contrast, interactions among glial cells, myocytes, and stromal cells, particularly glial cell-myocytes was reduced in pouch failure tissues relative to non-failure tissues.

In conclusion, TAC specimen samples from patients with subsequent pouch failure showed altered cellular neighborhoods characterized by dense immune cell hubs and fewer interactions between cells that maintain gut motility, neural activity, and structural support (i.e., myocytes, glial cells, and stromal cells).

### Spatial gene set enrichment analysis identifies cytokine crosstalks associated with a high risk of pouch failure

To identify the molecules involved in the pathogenic cellular interactions, we performed a spatial gene set enrichment analysis. In the first step, differential gene expression analysis was performed between pouch failure and non-failure samples of a single cell type within a given tissue layer. Epithelial cells showed a strong transcriptional signature of type I/II IFN in pouch failure samples with the overexpression of genes such as *OAS3, MX1, OAS1, OASL* (FC > 10, *P* < 0.03, **Figure 5A, B).** Macrophages, central to the pouch failure phenotype, exhibited distinct expression patterns depending on the tissue layer. In the mucosa, they overexpressed inflammatory molecules such as *CXCL5, CXCL8, IDO1, S100A8, IL-1β, OASL,* and *IL6* (**Figure 5C)**. In the muscularis propria, they had a different expression profile, which included *S100A8, S100A9, STAT1, IL1R2, and IL3RA* (**Figure 5F)**.

**Figure 5.**
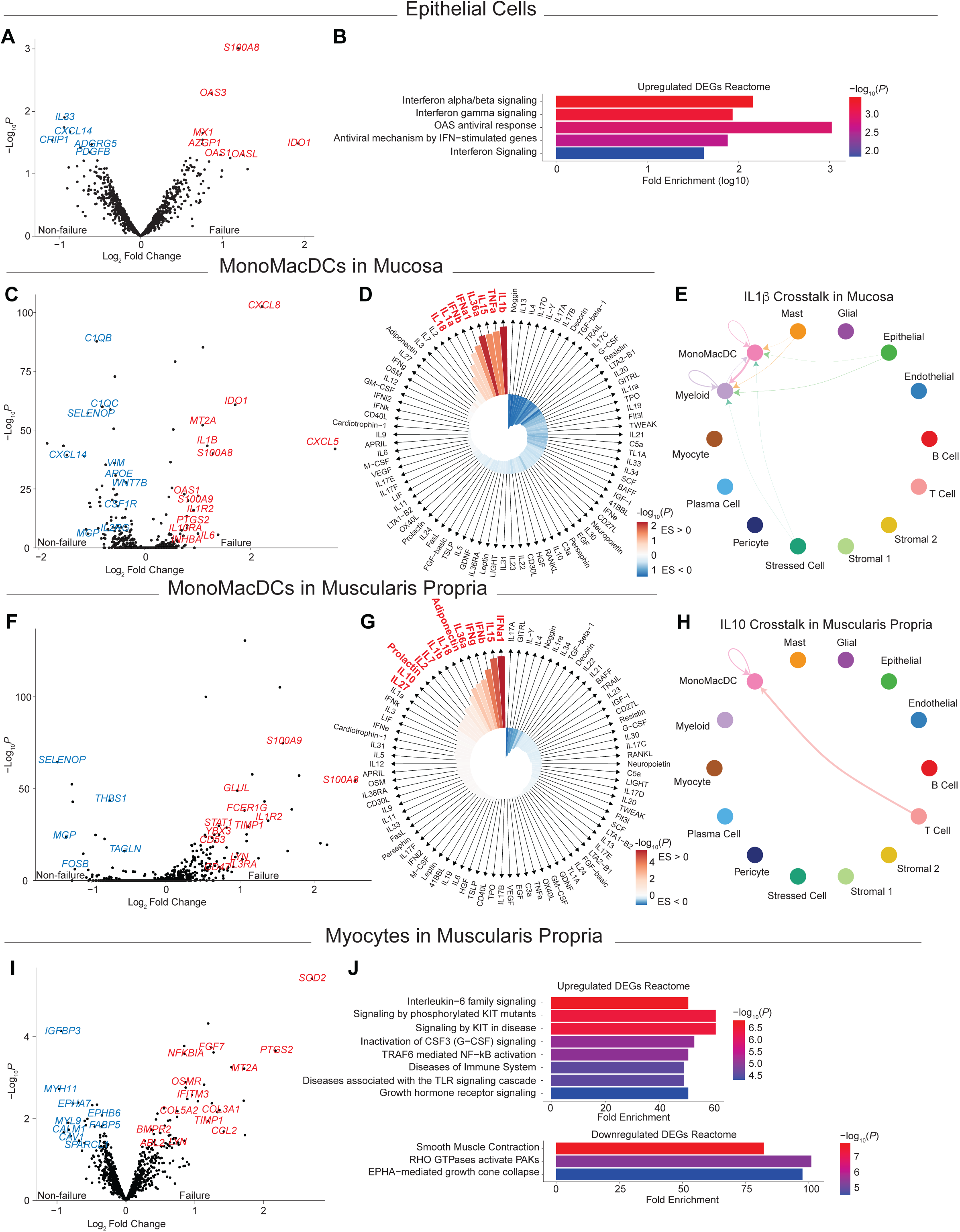
Spatial gene expression and cytokine signature enrichment analysis reveal type I/II IFN, IL-1β, and IL-10-driven cellular interactions linked to increased risk of future pouch failure. **(A)** Volcano plot showing differential gene expression in epithelial cells from failure vs. non-failure patients. Upregulated and downregulated genes in red and blue, respectively. *P-values* from moderated two-tailed t-test. **(B)** Gene set enrichment analysis of upregulated genes from (A) using the Reactome database. Fold enrichment on the x-axis. Color intensity proportional to the Bonferroni-adjusted *P-value* (one-sided hypergeometric test). **(C)** Volcano plot similar to (A) showing differential gene expression in MonoMacDCs in the mucosa from failure vs. non-failure patients. **(D)** IREA cytokine enrichment plot^37^ showing enrichment scores for 86 cytokine signatures in MonoMacDC within the mucosa of failure vs. non-failure patients. Bar height represents the enrichment score (ES), and shading denotes the FDR-adjusted *P*-*value* (hypergeometric test), with darker shades indicating higher significance. Red and blue show enrichment in failure and non-failure, respectively. **(E)** Circle plot representing the IL-1β-mediated cellular crosstalk network inferred by CellChat in the mucosa.^36^ Arrows indicate the direction from sender to receiver cells, with width proportional to the inferred interaction strength. **(F)** Volcano plot similar to (A) showing differential gene expression in MonoMacDCs in the muscularis propria from failure vs. non-failure patients. **(G)** IREA cytokine enrichment plot^37^ like (D) but in MonoMacDCs within the muscularis propria of failure vs. non-failure patients. **(H)** Circle plot like (E) but representing the IL-10-mediated cellular crosstalk network inferred by CellChat in the muscularis propria.^36^ **(I)** Volcano plot similar to (A) showing differential gene expression in myocytes from the muscularis propria in failure vs. non-failure patients. **(J)** Gene set enrichment analysis like (B) but of upregulated and downregulated genes from (I) using the Reactome database. Fold enrichment is displayed on the x-axis, with color intensity proportional to the Bonferroni-adjusted *P-value* (one-sided hypergeometric test).

To understand the alterations of immune signaling pathways in the pouch failure tissue, we then compared these responses with systematic studies of the gene signatures induced by cytokines in immune cells (Immune Dictionary)^37^. Geneset enrichment analysis showed that macrophage gene expression in failure samples within the mucosa and the muscularis propria were enriched for inflammatory cytokine responses, albeit with different patterns (**Figure 5D, G)**. While TNFα, IFNα1, IFNβ, IL-1β, IL-15, IL-18, and IL-36α signature genes were upregulated in both mucosa and muscularis propria, IL-1α response genes were selectively upregulated in the mucosa (**Figure 5D**) and IL-2, IL-7, IFNγ responses were upregulated in the muscularis propria (**Figure 5G)**. Genes induced by the anti-inflammatory cytokine IL10 were only upregulated in the muscularis propria (**Figure 5G)**.

To identify the cellular hubs associated with IL-1β, the most upregulated response in the mucosa, and IL-10, an inhibitory cytokine, in the muscularis propria, we performed a cellular proximity analysis of ligand and receptor-expressing cells (within 50 µm) using CellChat^36^. In the mucosa, MonoMacDC and myeloid cells were the main senders and receivers of IL-1β, with contributions from epithelial cells, mast cells, and stressed cells (**Figure 5E)**. In the muscularis propria, IL-10 was mainly produced by T cells to affect MonoMacDCs (**Figure 5H)**.

Myocytes also showed a specific gene expression signature in pouch failure samples. Myocytes overexpressed inflammatory genes (e.g., *IFITM3, CCL2,* and *OSMR*), enriched with IL6 family signaling, TLR signaling, and downregulated genes associated with muscle contraction (e.g., *MYH11*, *MYL9*, and *CALM*) (**Figure 5I, J)**.

In conclusion, spatial analysis of gene expression programs and ligand-receptor expression identified a role for type I/II IFN in the epithelium, IL-1β in the mucosa, and IL-10 pathways in the muscularis propria of colectomy specimens from patients with subsequent pouch failure. These cytokines involved specific cellular interactions between immune (in particular MonoMacDC and T cells) and non-immune cells (epithelial cells and myocytes).

## Discussion

This study identified TAC histologic features linked to a high risk of pouch failure in IPAA patients and utilized spatial transcriptomics to explore the underlying biological processes of this susceptibility. Systematic histologic assessment of inflammation severity and depth revealed key indicators of pouch failure risk, notably the depth of immune cell infiltration.

Spatial transcriptomics revealed that the infiltration was primarily composed of MonoMacDC and T cells within the muscularis propria and suggested potentially disrupted interactions between myocytes and glial cells. Lastly, we identified targetable cytokine pathways that could inform patient management and enhance IPAA outcomes. Our findings highlight the importance of preoperative assessment, detailed histologic evaluation, and molecular profiling to identify patients predisposed to pouch failure.

This study followed a longitudinal University of Chicago cohort for up to 18 years. The 10% failure rate matched national averages for pouch failure^5–7^, and, when accounting for loss to follow-up, the estimated risk of pouch failure rose to 23%, stressing the importance of better selecting IPAA candidates^8^. However, few studies have examined the link between TAC histologic features (pre pouch formation) and IPAA outcomes beyond the traditional pathological evaluation of TAC specimens, typically concluding with a diagnosis of UC, CD, or IC. Tan et al. ^40^ reported that chronic inflammation of the muscularis propria was linked to pouch-related complications, while Yantiss et al.^41^ identified fissuring ulcers as a predictor of pouchitis^39, 40^. To better identify patients at risk of pouch failure, we propose a structured histologic evaluation system that assesses TAC specimens every 10 cm, focusing on the depth of chronic inflammation, disease activity severity, and high-risk features of CD. The presence of at least one high-risk histologic feature was the strongest predictor of pouch failure, increasing the risk ninefold. IPAA failure patients had larger areas of severe active disease, deeper chronic inflammatory infiltrates (especially within the muscularis propria), and more frequent “knife-like” ulcers extending into the deep submucosa or muscularis propria.

In managing IBD patients, our TAC evaluation system with a synoptic reporting structure offers a new approach by providing more comprehensive and clinically relevant information than pre-colectomy biopsies, which often miss critical high-risk features due to limited and superficial tissue sampling. It also precedes the Chicago Classification, which assesses patients only after pouch formation. Like in the TAC, inflammation remains a key predictor of pouch failure by endoscopy (inlet stenosis, diffuse inflammation, and cuffitis)^17^. Notably, TAC histologic findings correlated with endoscopic evidence of pouch inflammation.

The “conversion” to CD or “de novo” CD of the pouch following an initial diagnosis of UC or IC remains a controversial concept. Reclassification to CD of the pouch develops in ∼13% of patients (range: 4-30%)^10, 11, 14, 41–43^ and our rate of 22% (62/280) aligns with findings from large referral centers. However, it remains unclear whether these patients truly develop CD over time, or if they had CD limited to the colon that later manifests in the small bowel post-pouch formation. Understanding this progression is clinically relevant, as it could guide preoperative counseling and postoperative management, especially for patients with high-risk histologic features in their TAC specimens. We proposed a nuanced classification of IPAA with Crohn’s-like features^8^ and the term “pouch with Crohn’s disease-like features” instead of “Crohn’s disease of the pouch” for patients who develop Crohn’s-like inflammation after IPAA without a pre-colectomy diagnosis of Crohn’s disease. Through endoscopic evaluation of the pouch, we previously identified six phenotypes of pouches with Crohn’s disease-like features^17^, each associated with distinct risks of pouch failure and complications. This revised terminology more accurately reflects the phenotypic overlap and diagnostic complexity of these cases, acknowledging a spectrum of disease presentations that may not align with a traditional CD of the pouch diagnosis^8^. Incorporating systematic histologic classification of the TAC, as demonstrated in this study, could enhance diagnostic precision and support tailored therapeutic interventions through more objective, reproducible measures.

Spatial transcriptomics provided a powerful tool to link histologic findings with underlying cellular and molecular mechanisms. Changes in the epithelium were associated with a stress-related transcriptional response to type I and II interferons (IFNs). The importance of the depth of inflammation was also evident through spatial transcriptomics, which further identified the infiltrating cell types, specifically macrophages and T cells. These cells penetrated deep into the muscularis propria of failure patients and displayed distinct molecular signatures based on tissue location and risk of failure. In the mucosa, macrophages in failure patients expressed genes responsive to pro-inflammatory cytokines such as TNFα, IFNγ, and IL-1β, produced by immune, stromal, and epithelial cells. In contrast, macrophages deeper in the muscularis propria expressed genes related to the anti-inflammatory cytokine IL-10. The roles of these cytokine signaling pathways in IBD are well documented: IL-10 is linked to immune regulation and tissue repair, while IFN, TNFα, and IL-1β signaling typically promote inflammation and immune activation^37^. Although the presence of both pro– and anti-inflammatory signals may seem contradictory, the spatial context reveals that these signals operate in distinct tissue locations. Our findings also suggest that non-immune cell dysfunction in the muscularis propria may contribute to adverse outcomes. We observed disrupted communication between glial cells and myocytes within the muscularis propria, and muscle cells showed reduced expression of contraction-related genes, such as myosin. This disruption of physiological interactions may impair gut motility, potentially leading to pouch dysfunction and failure.

This analysis offers potential options for therapy and IPAA risk stratification. Anti-TNF therapy, for example, is already a standard treatment for IBD^44^. For future molecular diagnostics, immunohistochemistry staining or spatial transcriptomics studies of T cells (e.g., CD3 IHC) and macrophages (e.g., CD68 IHC) in the TAC could help quantify the depth and extent of infiltration for risk stratification. Understanding the roles of glial cells and myocytes in IBD and IPAA failure may also open new therapeutic avenues, including strategies to preserve or restore their critical cellular interactions.

In conclusion, our study advances the understanding of IPAA failure, enhances clinical management strategies, and reveals new molecular pathways for targeted interventions. By implementing a structured histological assessment focused on high-risk histologic features and spatial molecular analysis, patients at risk can be more effectively stratified, guiding both surgical decisions and postoperative care. This work exemplifies how integrating clinical, histological, and systems biology approaches can foster a transformative area of translational research.

## Author contributions

ADO, KU, DZ, and CRW conceptualized the study. ADO and CRW performed the histological analyses. PCMN, ES, and DZ conducted the spatial transcriptomics analyses. JFC carried out statistical analyses. ADO, PCMN, ES, DZ, and CRW wrote the initial manuscript. LS, KSO, DR, EBC, and NH critically reviewed and revised the manuscript. ADO, CRW, SA, DR, NH, KU, and KSO provided clinical and surgical expertise and curated patient samples. All authors reviewed and approved the final manuscript.

## Data, analytic methods, and study materials availability

Spatial transcriptomics data is available on Gene Expression Omnibus (Study GSE283625, reviewer token: obsnykogbpipdcv). Bioinformatic code is published on GitHub (website https://github.com/zemmourlab/pouch_git). Requests for study materials will be considered after discussion with the corresponding authors.

## References

1. Lovegrove RE, Heriot AG, Constantinides V, et al. Meta-analysis of short-term and long-term outcomes of J, W and S ileal reservoirs for restorative proctocolectomy. Colorectal Dis 2007;9:310–20.

2. Nicholls RJ, Pezim ME. Restorative proctocolectomy with ileal reservoir for ulcerative colitis and familial adenomatous polyposis: a comparison of three reservoir designs. Br J Surg 1985;72:470–4.

3. Gu J, Remzi FH, Shen B, et al. Operative strategy modifies risk of pouch-related outcomes in patients with ulcerative colitis on preoperative anti-tumor necrosis factor-alpha therapy. Dis Colon Rectum 2013;56:1243–52.

4. Pandey S, Luther G, Umanskiy K, et al. Minimally invasive pouch surgery for ulcerative colitis: is there a benefit in staging? Dis Colon Rectum 2011;54:306–10.

5. Hueting WE, Buskens E, van der Tweel I, et al. Results and complications after ileal pouch anal anastomosis: a meta-analysis of 43 observational studies comprising 9,317 patients. Dig Surg 2005;22:69–79.

6. Mark-Christensen A, Erichsen R, Brandsborg S, et al. Pouch failures following ileal pouch-anal anastomosis for ulcerative colitis. Colorectal Dis 2018;20:44–52.

7. Meagher AP, Farouk R, Dozois RR, et al. J ileal pouch-anal anastomosis for chronic ulcerative colitis: complications and long-term outcome in 1310 patients. Br J Surg 1998;85:800–3.

8. Akiyama S, Dyer EC, Rubin DT. Diagnostic and Management Considerations for the IPAA With Crohn’s Disease-Like Features. Dis Colon Rectum 2022;65:S77–S84.

9. Braveman JM, Schoetz DJ, Jr., Marcello PW, et al. The fate of the ileal pouch in patients developing Crohn’s disease. Dis Colon Rectum 2004;47:1613–9.

10. Brown CJ, Maclean AR, Cohen Z, et al. Crohn’s disease and indeterminate colitis and the ileal pouch-anal anastomosis: outcomes and patterns of failure. Dis Colon Rectum 2005;48:1542–9.

11. Hercun J, Cote-Daigneault J, Lahaie RG, et al. Crohn’s Disease after Proctocolectomy and IPAA for Ulcerative Colitis. Dis Colon Rectum 2021;64:217–224.

12. Melton GB, Fazio VW, Kiran RP, et al. Long-term outcomes with ileal pouch-anal anastomosis and Crohn’s disease: pouch retention and implications of delayed diagnosis. Ann Surg 2008;248:608–16.

13. Reese GE, Lovegrove RE, Tilney HS, et al. The effect of Crohn’s disease on outcomes after restorative proctocolectomy. Dis Colon Rectum 2007;50:239–50.

14. Shen B, Fazio VW, Remzi FH, et al. Risk factors for clinical phenotypes of Crohn’s disease of the ileal pouch. Am J Gastroenterol 2006;101:2760–8.

15. Tekkis PP, Heriot AG, Smith O, et al. Long-term outcomes of restorative proctocolectomy for Crohn’s disease and indeterminate colitis. Colorectal Dis 2005;7:218–23.

16. Yu CS, Pemberton JH, Larson D. Ileal pouch-anal anastomosis in patients with indeterminate colitis: long-term results. Dis Colon Rectum 2000;43:1487–96.

17. Akiyama S, Ollech JE, Rai V, et al. Endoscopic Phenotype of the J Pouch in Patients With Inflammatory Bowel Disease: A New Classification for Pouch Outcomes. Clin Gastroenterol Hepatol 2022;20:293–302 e9.

18. Fazio VW, Tekkis PP, Remzi F, et al. Quantification of risk for pouch failure after ileal pouch anal anastomosis surgery. Ann Surg 2003;238:605–14; discussion 614-7.

19. Frese JP, Grone J, Lauscher JC, et al. Risk factors for failure of ileal pouch-anal anastomosis in patients with refractory ulcerative colitis. Surgery 2022;171:299–304.

20. Shen B. Pouchitis: pathophysiology and management. Nat Rev Gastroenterol Hepatol 2024;21:463–476.

21. de Silva HJ, Jones M, Prince C, et al. Lymphocyte and macrophage subpopulations in pelvic ileal pouches. Gut 1991;32:1160–5.

22. Stallmach A, Schafer F, Hoffmann S, et al. Increased state of activation of CD4 positive T cells and elevated interferon gamma production in pouchitis. Gut 1998;43:499–505.

23. Goldberg PA, Herbst F, Beckett CG, et al. Leucocyte typing, cytokine expression, and epithelial turnover in the ileal pouch in patients with ulcerative colitis and familial adenomatous polyposis. Gut 1996;38:549–53.

24. Patel RT, Bain I, Youngs D, et al. Cytokine production in pouchitis is similar to that in ulcerative colitis. Dis Colon Rectum 1995;38:831–7.

25. He S, Bhatt R, Brown C, et al. High-plex imaging of RNA and proteins at subcellular resolution in fixed tissue by spatial molecular imaging. Nat Biotechnol 2022;40:1794–1806.

26. Cole SR, Hernan MA. Adjusted survival curves with inverse probability weights. Comput Methods Programs Biomed 2004;75:45–9.

27. Gramlich T, Petras RE. Pathology of inflammatory bowel disease. Semin Pediatr Surg 2007;16:154–63.

28. Stringer C, Wang T, Michaelos M, et al. Cellpose: a generalist algorithm for cellular segmentation. Nat Methods 2021;18:100–106.

29. Lopez R, Regier J, Cole MB, et al. Deep generative modeling for single-cell transcriptomics. Nat Methods 2018;15:1053–1058.

30. Martin PCN, Kim H, Lovkvist C, et al. Vesalius: high-resolution in silico anatomization of spatial transcriptomic data using image analysis. Mol Syst Biol 2022;18:e11080.

31. Hao Y, Stuart T, Kowalski MH, et al. Dictionary learning for integrative, multimodal and scalable single-cell analysis. Nat Biotechnol 2024;42:293–304.

32. Ritchie ME, Phipson B, Wu D, et al. limma powers differential expression analyses for RNA-sequencing and microarray studies. Nucleic Acids Res 2015;43:e47.

33. Chen Y, Chen L, Lun ATL, et al. edgeR 4.0: powerful differential analysis of sequencing data with expanded functionality and improved support for small counts and larger datasets. bioRxiv, 2024.

34. Pateiro-Lopez B, Rodriguez-Casal A. alphahull: Generalization of the Convex Hull of a Sample of Points in the Plane. 2.2 ed: The Comprehensive R Archive Network (CRAN), 2022.

35. Milacic M, Beavers D, Conley P, et al. The Reactome Pathway Knowledgebase 2024. Nucleic Acids Res 2024;52:D672–D678.

36. Jin S, Plikus MV, Nie Q. CellChat for systematic analysis of cell-cell communication from single-cell transcriptomics. Nat Protoc 2024.

37. Cui A, Huang T, Li S, et al. Dictionary of immune responses to cytokines at single-cell resolution. Nature 2024;625:377–384.

38. Garrido-Trigo A, Corraliza AM, Veny M, et al. Macrophage and neutrophil heterogeneity at single-cell spatial resolution in human inflammatory bowel disease. Nat Commun 2023;14:4506.

39. Tan KK, Ravindran P, Young CJ, et al. The extent of inflammation is a predictor for pouch-related complications in ileal pouches in patients with ulcerative or indeterminate colitis. Colorectal Dis 2014;16:620–5.

40. Yantiss RK, Sapp HL, Farraye FA, et al. Histologic predictors of pouchitis in patients with chronic ulcerative colitis. Am J Surg Pathol 2004;28:999–1006.

41. Barnes EL, Kochar B, Jessup HR, et al. The Incidence and Definition of Crohn’s Disease of the Pouch: A Systematic Review and Meta-analysis. Inflamm Bowel Dis 2019;25:1474–1480.

42. Lightner AL, Fletcher JG, Pemberton JH, et al. Crohn’s Disease of the Pouch: A True Diagnosis or an Oversubscribed Diagnosis of Exclusion? Dis Colon Rectum 2017;60:1201–1208.

43. Lightner AL, Mathis KL, Dozois EJ, et al. Results at Up to 30 Years After Ileal Pouch-Anal Anastomosis for Chronic Ulcerative Colitis. Inflamm Bowel Dis 2017;23:781–790.

44. Gros B, Kaplan GG. Ulcerative Colitis in Adults: A Review. JAMA 2023;330:951–965.

