## Supplementary Methods and Figures for "Histopathologic Evaluation and Single-Cell Spatial Transcriptomics of the Colon Reveal Cellular and Molecular Abnormalities Linked to J-Pouch Failure in Patients with Inflammatory Bowel Disease"

*Study Cohort*

Patients with a diagnosis of IBD who underwent IPAA from 2000-2010 at the University of Chicago Medical Center were identified from a prospectively maintained, IRB-approved surgical database (IRB 16014A). The primary clinical outcome was categorized as either IPAA failure or IPAA non-failure. IPAA failure was defined as those patients requiring pouch excision or permanent diverting ileostomy. Electronic medical records were retrospectively reviewed to determine pouch duration and the nature of pouch failure, *i.e.,* “medical failure” versus “technical failure.” Cases determined to be technical failures (early surgical complications occurring < 3 months from IPAA) and one patient with complications from metastatic cancer were excluded from the study (n=3).

*Definition of Crohn’s Disease*

In this study, the term "Crohn's disease (CD)" is used broadly to encompass all categories along its clinical diagnostic continuum, including (1) pre-TAC diagnosis of Crohn’s disease, (2) CD diagnosed histologically in total abdominal colectomy specimen, and (3) cases initially diagnosed as ulcerative or indeterminate colitis that evolve into a CD phenotype post-TAC or IPAA. The latter includes “pouchitis with CD-like features,” and patients who develop CD-like strictures of the pre pouch ileum. Unless otherwise specified, we did not subcategorize patients according to the time of diagnosis.

*Pouch Survival Analysis*

Survival analysis of the entire cohort (n=414) was conducted to compare the time to pouch failure between patients subsequently diagnosed with CD of the pouch and those with non-CD pouch diagnoses. Kaplan-Meier survival curves were generated to estimate the cumulative probability of pouch survival over time in each group. Time was measured from the date of IPAA surgery to the date of pouch failure or last follow-up. A log-rank test was performed to assess statistical significance between the survival distributions of the two groups. Cox proportional hazards and hazard ratios (HR) with corresponding 95% confidence intervals (CI) were calculated to quantify the relative risk of pouch failure in patients with CD versus non-CD diagnoses. Inverse probability weighting (IPW) Kaplan-Meier analysis was used to estimate survival probabilities based on inflammation severity and depth. The weights were derived from the inverse probability of group membership to adjust survival estimates, addressing sampling bias and ensuring comparability between groups.[26] All analyses were conducted using R with the survival v3.7-0[45] and survminer v0.5.0 packages.[46] Statistical significance was set at $P < 0.05$.

*Histologic Evaluation*

Histologic analysis of TAC specimens was performed on a subset of the cohort. Of the 41 patients with IPAA failure, slides from 39 patients were available for review. These patients were matched 1:1 for age, sex, and number of operative stages with IPAA non-failure patients who reported satisfactory pouch function for at least 2 years (case-control cohort). Additional clinicopathologic information recorded for all patients included: disease duration, use of preoperative biologics, preoperative treatment of *C. difficile* colitis, preoperative biopsy diagnosis, indication for TAC, TAC pathologic diagnosis, and post-IPAA follow-up time.

Hematoxylin and eosin (H&E)-stained sections of TAC specimens, which included the proximal (ileal) and distal resection margins, appendix, and sampling of the entire specimen every 10 cm from proximal to distal, were independently reviewed in a blinded fashion by two gastrointestinal pathologists. The severity of disease activity throughout each specimen was graded according to previously published criteria[27] **(Table S1)** and the overall average percentage of ulceration (severe disease activity) was calculated. The depth of chronic inflammatory infiltrates throughout each specimen was assigned a score 0-4 in the following manner: score 0- no increase in chronic inflammation (normal), score 1- increased chronic inflammation within the mucosa only, score 2- increased chronic inflammation within mucosa and extension into submucosa, score 3- chronic inflammation extending from submucosa into muscularis propria, and score 4- chronic inflammation extending from submucosa through muscularis propria and into subserosal/pericolonic fibroadipose tissue (transmural inflammation) **(Table S2)**. "Non-UC" histologic features were also recorded, including the presence of epithelioid granulomas (not associated with crypt rupture), active ileal inflammation with evidence of chronic injury, deep penetrating "knife-like" ulcers (distinct ulcers extending to the deep submucosa/muscularis propria), right-sided disease with rectal sparing, and patchy non-continuous colonic disease involvement. Histologic findings were then correlated with clinical pouch outcome.

*Stepwise Logistic Regression*

Stepwise logistic regression was performed using analysis of maximum likelihood estimates to determine which histologic features of the TAC specimen along with *preoperative* patient-specific characteristics of the case-control cohort were associated with pouch outcome. Clinical

characteristics included in the logistic regression model were as follows: matched variables (age, sex, number of operative stages), duration of disease, preoperative diagnosis on biopsy, use of preoperative biologics, preoperative treatment of *C. difficile* colitis, and indication for colectomy. Histologic features of the TAC specimen included in the logistic regression model were the average percentage of ulceration and the presence of one or more of the following histologic features: depth of inflammation score ≥ 3 (in at least one histologic section), epithelioid granulomas not associated with crypt rupture, ileal inflammation with evidence of chronic injury, deep penetrating knife-like ulcers, right-sided disease, and non-continuous colonic disease involvement. The *c*-statistic was used to estimate the goodness of fit of the model.
Logistic regression was performed to evaluate the association between histological scores (inflammation depth and severity) and the Chicago Classification of pouchoscopy.[17] The glm(family = binomial(link='logit')) function was used from the stats v4.4.2 in R.[47]

*Spatial Transcriptomics Using Bruker's CosMx Spatial Molecular Imager*

The CosMx Spatial Molecular Imager (SMI) was used for spatial transcriptomics. Tissue samples analyzed were from the left colon of TAC specimens (4 pouch failure patients and 4 non-failure patients; 1 section per patient). Formalin-fixed paraffin-embedded (FFPE) tissue sections were cut at 5 µm thickness and mounted on Leica BOND plus slides. Following the manufacturer's instructions (Manual Slide Preparation, MAN-10159-01, November 2022), sections were hybridized with the 1K Human Universal Cell Characterization Panel, comprising 974 spatially barcoded probes targeting 950 genes of interest, 10 control genes and a 20-gene custom panel (CCR4, CCR6, CD57, CD73, EBI3, EGR1, FOSB, GPR15, GPR25, HOPX, ID2, IKZF2, KLRG1, LEF1, NR4A1, RORC, SLC2A3, TGFBI, TXNIP, ZFP36L2). Cell segmentation was aided by the Universal Segmentation Kit (DAPI, CD298/B2M), Supplemental Segmentation Kits (PanCK, CD45), and a CD3 marker. On Day 0, sections were baked overnight at 60 °C. On Day 1, after deparaffinization and rehydration, antigen retrieval was performed by immersing the slides in CosMx Target Retrieval Solution and heating them in a pressure cooker at 110 °C for 15 minutes. Subsequently, the slides are treated with digestion buffer and fiducials were added for precise imaging alignment. Hybridization of the CosMx RNA probe was performed at 37°C overnight in a hybridization oven. On day 2, nuclear and cell segmentation staining was performed for 1h at room temperature in the dark. The CosMx flow cell was then assembled onto the slide, creating a sealed environment that prevents tissue drying and facilitates controlled reagent flow for high-resolution imaging on the CosMx SMI platform. On average, 35.5 fields of view (FoVs) were captured per sample, resulting in a total of 284 FoVs. The mean area examined per sample was 9.10 $mm^2$. RNA transcript readout on the CosMx SMI instrument was performed as previous described.[25] Briefly, RNA readout began with the sequential addition of reporter probe pools, each followed by imaging, UV-cleavage of fluorophores, and washing. This process was repeated across 16 reporter pools.

*Cell Segmentation and Quality Control*

Following the manufacturer's instructions for the bioinformatic platform Atomx (CosMx SMI Instrument User Manual, MAN-10161-01, November 2022), cells were segmented using CellPose[28] with the default configuration A to define cell boundaries, assign transcripts at the single-cell level, and generate a transcript-by-cell count matrix. The gene panel also includes 10 External RNA Controls Consortium (ERCC) negative target probes containing hybridization regions not complementary to the human transcriptome. Non-specific probe hybridization was quantified by the mean count of negative control probes per probe per cell. Additionally, barcode reading errors were quantified as the mean false code fraction per cell, i.e. the fraction of false codes detected relative to the transcript count. Cells were filtered out if they had more than 5% negative control probe counts, or fewer than 10 transcript counts (Supplementary Table 3). Additionally, cell area outliers were identified and removed using the interquartile range (IQR) method, with cells having an area more than 1.5 times the IQR below the first quartile or above the third quartile being excluded. The IQR method was also applied to filter out FoVs where the mean count per cell was 1.5 times the IQR below the first quartile. After applying these quality control measures, 96.53% of cells across samples and FoVs were retained.

*Preprocessing, Integration, and Dimensionality Reduction*

Data processing was primarily conducted using the Seurat library (V5.1.0)[31] with the default settings for NormalizeData, and ScaleData applied to all genes. SCVI (V1.0.0)[29] was used to correct for batch effects across patient samples and FoVs. In the SCVI model, all genes were utilized, with the batch_key set to sample_id and the categorical_covariate_key set to FoV. The SCVI model was trained using parameters of n_layers = 2, n_latent = 30, and gene_likelihood = "nb". All 30 dimensions of the SCVI latent space were used as the input for FindNeighbors() with k.param = 15 and FindClusters() with a resolution of 1 in Seurat (V5.1.0). Uniform Manifold Approximation and Projection (UMAP) was performed using the 30-dimensional SCVI latent space via the RunUMAP function with min.dist = 0.01, spread = 5, and metric = "cosine." Three clusters ("9", "19", and "20") were primarily located within the blood vessels of the samples identified as blood cells and were excluded from downstream analysis.

*Cell Type Annotation*

To identify the top markers in each cluster, FindAllMarkers() in Seurat V5.1.0 was run with a logfc.threshold of 0.25, min.pct of 0.05, and base = 2. The top 10 genes, ranked by average log2FC per cluster, were visualized in a heatmap, and cell types were manually annotated based on known marker genes. T cells, B cells, and the Stromal 2 population, initially identified, were further subclustered to explore cell type subsets and the annotations were updated based on marker genes in the subclusters. A dot plot displayed selected known marker genes for each cell type.

*Tissue Layer Identification*

Vesalius (V2.0.0)[30] was used for tissue layer identification, with image embeddings generated through a 3-dimensional UMAP. Tiles were produced using tensor_resolution = 1, filter_grid = 0.01, and filter_threshold = 0.99. The images were regularized using the default settings, except for 20 iterations. Tissue segmentation into 8 regions was performed using the segment_image function with k-means clustering, allowing the identification of the complex-shaped epithelium layer. Manual annotation using tidygate (V1.0.14)[48] was applied to the epithelium-removed data to identify more regularly shaped layers, including the muscularis mucosae, submucosa, muscularis propria, and subserosa. The regularized, smoothed, and k-means segmented Vesalius images, alongside H&E-stained images, were used to guide the definition of these layer boundaries.

*Differentiation Expression Analysis*

Differential expression was analyzed between pouch failure and non-failure samples for each cell type per tissue layer. The analysis was performed using FindMarker() in Seurat V5.1.0 with the Wilcoxon Rank Sum test option or through a pseudobulk approach utilizing the limma (V3.58.1)[32] and EdgeR (V4.0.16).[33] In our limma-trend pipeline, library sizes were first normalized using the trimmed mean of *M-values* (TMM) normalization, followed by calculating log-transformed counts per million (CPM) with a prior count of 0.1. A linear model matrix based on J-pouch results was constructed and fit the normalized data. Empirical Bayes statistics with trend adjustment and robust fitting enabled were computed for differential expression. Significant genes ($P < 0.05$) were identified and visualized in volcano plots.

*Paired Nearest Neighborhood Analysis*

K Nearest Neighbors (KNN) is applied to cells in the muscularis propria with k=2 within a radius of 20 µm to identify cell pairs (target cells and their nearest neighbor within 20 µm). The EdgeR and limma-trend mentioned above were used to normalize the number of cell pairs per sample and to perform cell-pair differential analysis between pouch failure and non-failure samples.

*Minimum Distance Analysis*

In the minimum distance analysis, the Euclidian distance between each MonoMacDC cell and its closest neighboring cell of each type was calculated to determine the minimum distance. The average minimum distance per sample was then computed by averaging all the minimum distances for MonoMacDC-neighbor pairs within each sample. A t-test was performed to compare the average minimum distance for each MonoMacDC-neighbor pair between failure and non-failure samples.

*Cell Type Density Calculation*

The areas of the epithelium and lamina propria, which are densely filled with cells and have irregular shapes, were estimated by summing the total area of the cells within these layers for each sample. For the sparse and more regularly shaped layers (muscularis mucosae, submucosa, muscularis propria, and subserosa), the area was estimated using the alpha shape calculated with the alphahull library (V2.5).[34] The alpha shape was generated using the centroids of the cells as input, with alpha values optimized per sample layer, ranging from 0.05 to 0.3. The average global cellular density was then calculated by dividing the number of cells by the total area of the corresponding tissue layer on a per sample basis. Global cellular density was compared between pouch failure and non-failure samples using a one-sided Wilcoxon rank-sum test, and log2 fold changes were plotted to visualize the differences.

To obtain a more detailed view of cellular density distribution, local cellular density was calculated by counting the number of neighboring cells of the same cell type within a 100 μm x 100 μm window surrounding each target cell.

*Protein Data Processing and Visualization*

Protein expression data (CD3, CD45, β2m, PanCK) were normalized using a centered log-ratio transformation[49] across cells with the NormalizeData function in Seurat (V5.1.0). PanCK by CD45 and CD3 by CD45 were plotted for epithelial cells, immune cells (non-T cells), T cells, and other non-immune cells. To visualize the cell density in the protein expression plots, the kde2d() function from the MASS package (V7.3-60.0.1)[50] was used to highlight regions with high cellular density.

*scRNA-seq – Single-Cell Spatial Transcriptomics Comparison*

Ulcerative Colitis scRNA-seq dataset was obtained from the study by Alba Garrido-Trigo *et al*.[38] The *nanostring_reference* annotation provided by the authors was used, and cell types were grouped to match those in the CosMx dataset for comparison. The percentage of each cell type in both the scRNA-seq and CosMx datasets was plotted and compared.

*Pathway Enrichment Analysis*

Pathway enrichment analysis as performed on epithelial cells and myocytes, using up- and down-regulated genes from pseudobulk analysis as individual inputs. The run_pathfindR()[51] function from the pathfindR package (V2.3.1) was applied using the Reactome[35] gene sets with default settings, except for min_gset_size = 5.

*CellChat Analysis*

CellChat v2.0[36] and the Immune Dictionary App[37] were used to investigate intercellular communication. Differential Gene Expression was performed between failure and nonfailure MonoMacDCs in the mucosa and the muscularis propria. DE genes with a $P < 0.05$ were then interrogated for cytokine signatures using the Immune Dictionary App[37] and its Immune Response Enrichment Analysis module, focusing on macrophages as a cell type and utilizing the hypergeometric method for enrichment statistics. Cell Chat v2.0 was then used to identify the crosstalk in each layer involving IL-1 and IL-10. Focusing on "Secreted Signaling" pathways, CellChat objects were constructed for every group (by layer and by outcome), and communication probabilities were calculated based on differential “receptor” and “ligand” expressions using the computeCommunProb(). Gene names were truncated for compatibility. Interactions were filtered to only those in at least 100 cells. IL-1 and IL-10 circle plots were displayed using the netVisual_aggregate() function.

*Statistical Analysis*

Analysis was conducted using R-3.6.2. Univariate statistical analysis was performed using the Fisher Exact Test or 2-tailed Students *t*-test where appropriate. Additional statistical tests are described in their respective method section. *P-value*s or adjusted *P-values* using the Benjamini-Hochberg procedure of less than 5% were deemed significant.

**Supplementary References**


45. Therneau TM. A Package for Survival Analysis in R. 3.7-0 ed: The R Foundation, 2024.
46. Kassambara A, Kosinski M, Biecek P. survminer: Drawing Survival Curves using 'ggplot2'. 0.4.9 ed. Vienna, Austria: The R Foundation for Statistical Computing, 2023.
47. Bolar K. STAT: Interactive Document for Working with Basic Statistical Analysis: R package version 0.1.0, Comprehensive R Archive Network (CRAN), 2019.
48. Mangiola S, Jawaid W. tidygate: Interactively Gate Points (Version 1.0.14) [R package]: The Comprehensive R Archive Network (CRAN), 2024.
49. Aitchison J. The Statistical Analysis of Compositional Data. Journal of the Royal Statistical Society: Series B (Methodological) 1982;44:139–177.
50. Venables WN, Ripley BD. Modern Applied Statistics with S: Springer, 2002.
51. Ulgen E, Ozisik O, Sezerman OU. pathfindR: An R Package for Comprehensive Identification of Enriched Pathways in Omics Data Through Active Subnetworks. Front Genet 2019;10:858.

#### Figure S1

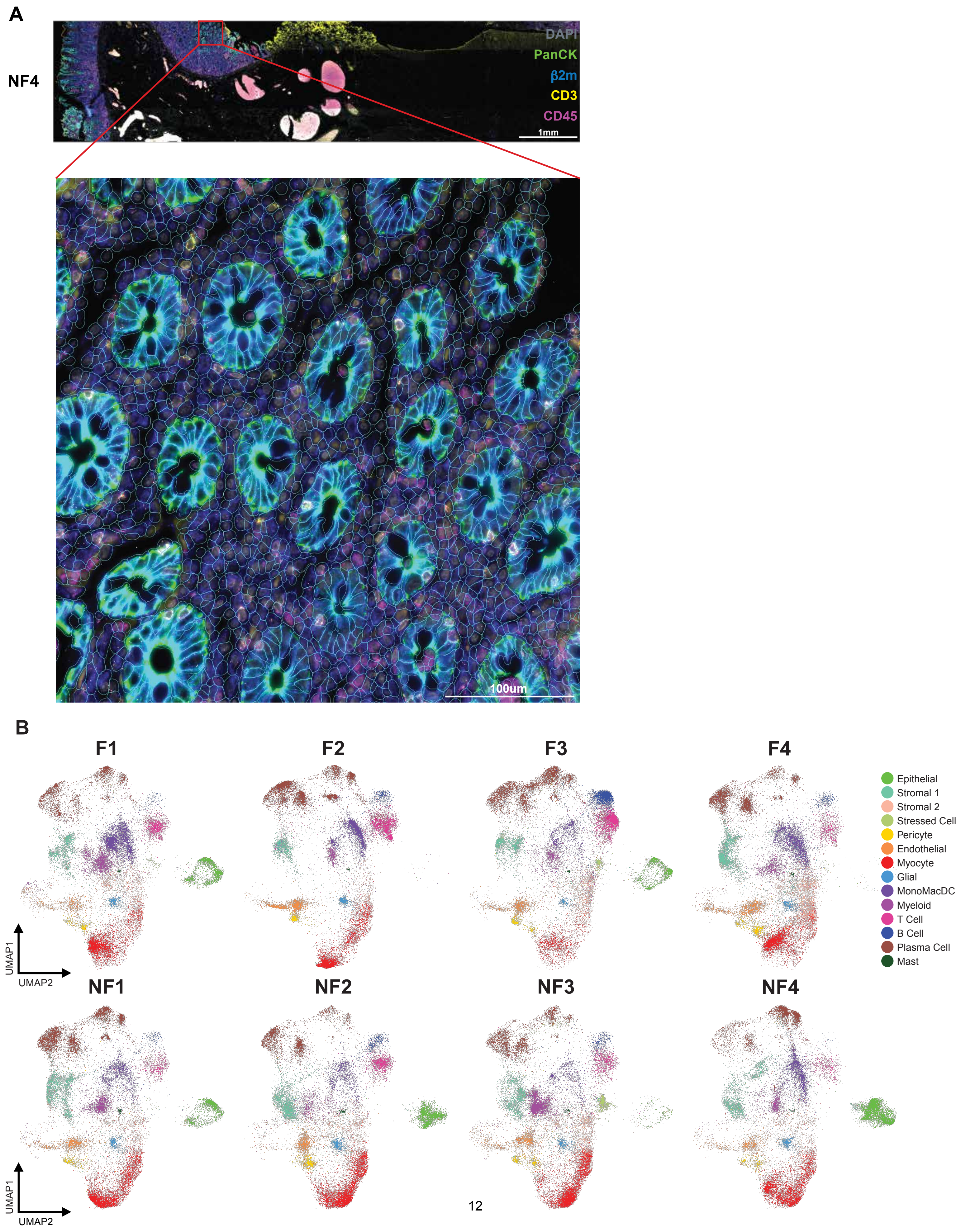
A
NF4
DAPI
PanCK
β2m
CD3
CD45
1mm
100um
B
F1
F2
F3
F4
NF1
NF2
NF3
NF4
UMAP1
UMAP2
Epithelial
Stromal 1
Stromal 2
Stressed Cell
Pericyte
Endothelial
Myocyte
Glial
MonoMacDC
Myeloid
T Cell
B Cell
Plasma Cell
Mast

#### Supplementary Legends

**Figure S1: Cell segmentation and batch correction evaluation in single-cell spatial transcriptomics (CosMx) samples**

(A) Cell segmentation using Cellpose (see Methods). The fluorescence image shows segmentation markers: DAPI (gray), PanCK (green), β2M (blue), CD3 (yellow), and CD45 (magenta). NF4 is shown as a representative sample, with an inset zoom into the mucosa as a representative example of the cell segmentation result (blue lines).

(B) 2D UMAP cell embedding after SCVI batch correction (for samples and FoVs, see Methods), colored by cell type and split by each sample, with an equal number of cells per sample shown (n = 28,428 per sample).

### Figure S2

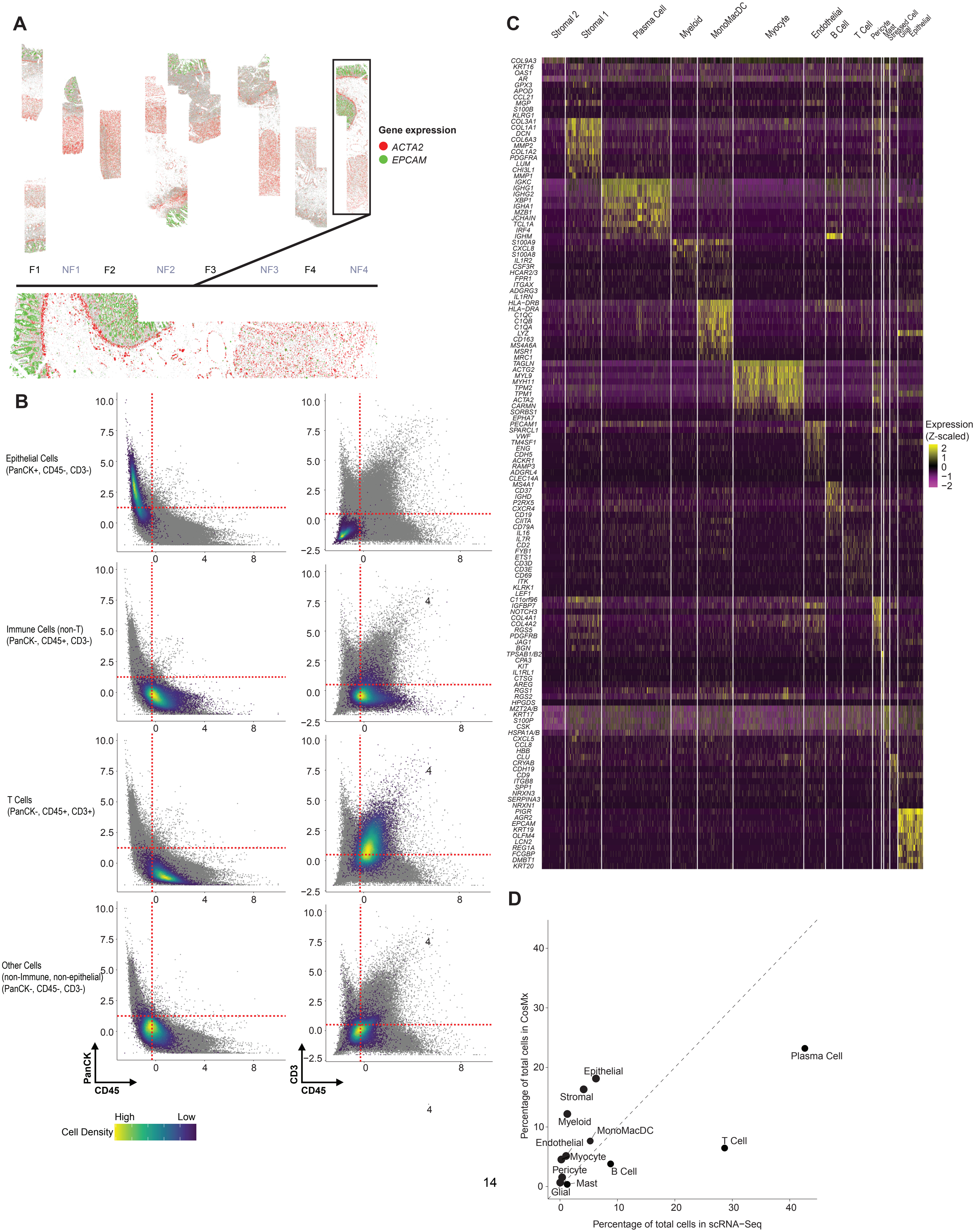

A
Gene expression
ACTA2
EPCAM
F1
NF1
F2
NF2
F3
NF3
F4
NF4
B
Epithelial Cells (PanCK+, CD45-, CD3-)
Immune Cells (non-T) (PanCK-, CD45+, CD3-)
T Cells (PanCK-, CD45+, CD3+)
Other Cells (non-Immune, non-epithelial) (PanCK-, CD45-, CD3-)
PanCK
CD45
CD3
CD45
High
Low
Cell Density
C
Stromal 2
Stromal 1
Plasma Cell
Myeloid
MonoMacDC
Myocyte
Endothelial
B Cell
T Cell
Pericyte
Mast
Stressed Cell
Glial
Epithelial
Expression (Z-scaled)
D
Percentage of total cells in CosMx
Percentage of total cells in scRNA−Seq
Plasma Cell
Epithelial
Stromal
Myeloid
MonoMacDC
Endothelial
T Cell
Myocyte
Pericyte
B Cell
Mast
Glial

**Figure S2: Unbiased identification of cell types and tissue layers by combining CosMx RNA and protein expression data.**

(A) Expression of *ACTA2* and *EPCAM* in colon samples shown in red and green, respectively, with dot size scaled to expression level. *EPCAM* marks the epithelial layer, while *ACTA2* highlights muscle-rich layers, including the muscularis mucosae and muscularis propria. The inset zooms into NF4 as a representative image, detailing the colon layers.

(B) Scatter plots showing normalized (Z-scaled and centered-log-ratio transformed) protein expression of segmentation markers PanCK, CD45, and CD3 in each single cell. Red dashed lines indicate thresholds separating positive and negative populations for each marker. Epithelial cells (PanCK+ CD45- CD3-), T cells (PanCK- CD45+ CD3+), other immune cells (PanCK- CD45+ CD3-), and other non-epithelial, non-immune cells (PanCK- CD45- CD3-) are color-coded on top of all other cells (in grey). The color gradient indicates their cell density in the plot.

(C) Heatmap displaying the Z-scaled expression of the top 10 marker genes for each cell type. Single cells are in columns, clustered by cell type.

(D) Scatter plot comparing cell type proportions between single-cell spatial transcriptomics CosMx data (this study) and reference single-cell RNA-seq data of ulcerative colitis from Alba Garrido-Trigo et al.[38] Note that this comparison is made on the mucosal layer, as the single-cell RNAseq dataset was generated from routine biopsies that do not contain deeper tissue layers.

### Figure S3

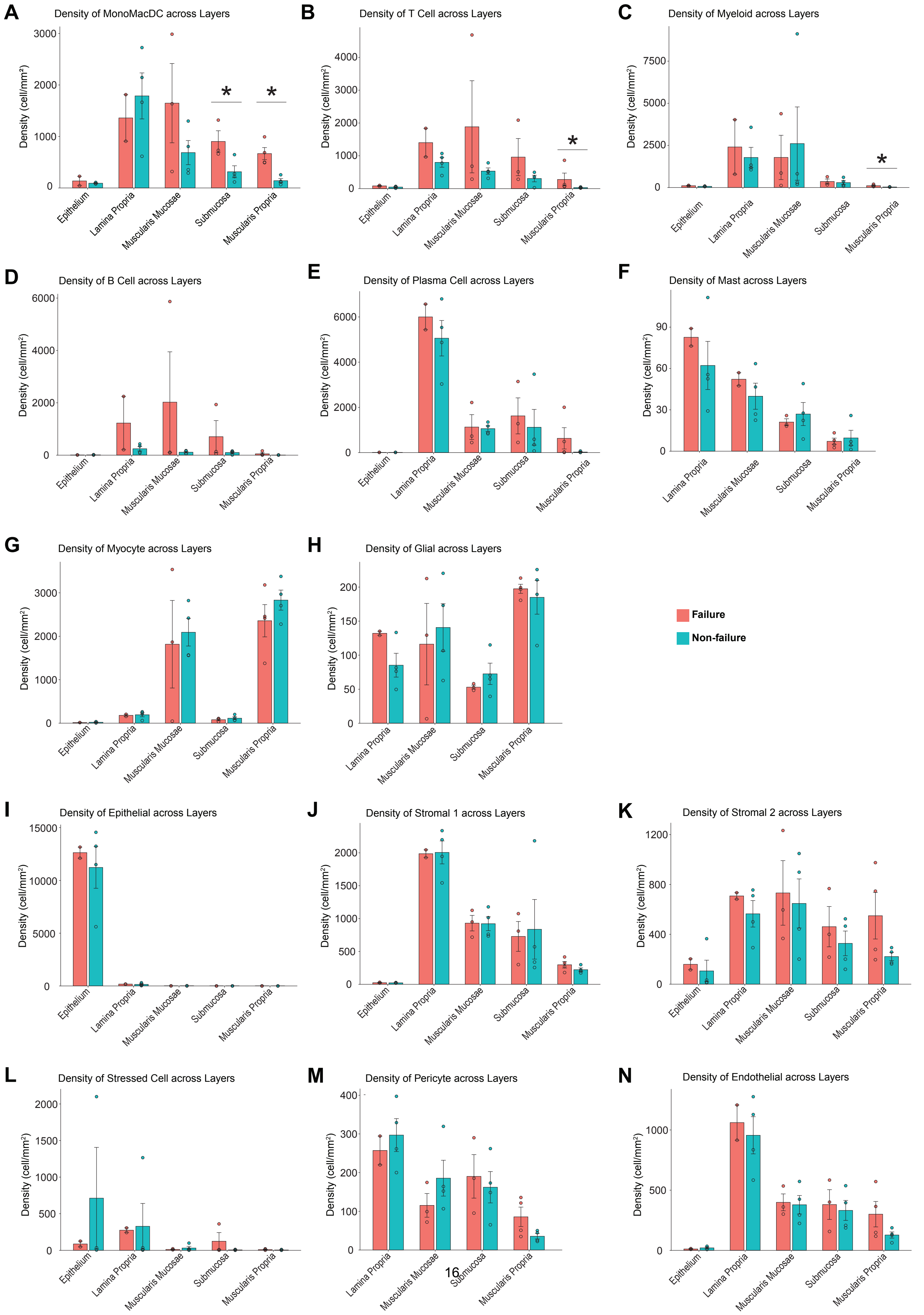

A
Density of MonoMacDC across Layers
B
Density of T Cell across Layers
C
Density of Myeloid across Layers
D
Density of B Cell across Layers
E
Density of Plasma Cell across Layers
F
Density of Mast across Layers
G
Density of Myocyte across Layers
H
Density of Glial across Layers
Failure
Non-failure
I
Density of Epithelial across Layers
J
Density of Stromal 1 across Layers
K
Density of Stromal 2 across Layers
L
Density of Stressed Cell across Layers
M
Density of Pericyte across Layers
N
Density of Endothelial across Layers
Density (cell/mm²)
Epithelium
Lamina Propria
Muscularis Mucosae
Submucosa
Muscularis Propria
*

**Figure S3: Comparative analysis of cell density across colon layers in failure vs. non-failure samples.**

(A–N) Barplots showing the cell density average and standard error of the mean (SEM) for immune and non-immune cell populations across the colon layers in failure vs. nonfailure samples (cells / $mm^2$). * $P < 0.05$, one-sided Wilcoxon rank-sum test.

**Table S1. Clinicopathologic characteristics in IPAA failure (n=39) and non-failure patients (n=39).**

Variables include demographic data, surgical staging, preoperative conditions, and pathologic diagnoses. Characteristics evaluated are disease duration, preoperative treatment, *C. difficile* infection, follow-up time, indications for TAC, and histopathologic diagnoses post-colectomy. *P-values* listed from Fisher's exact test

**Table S2. Histologic criteria for grading of disease activity.**

Mild activity is defined by the presence of neutrophils in the epithelium. Moderate activity is defined by neutrophils in the crypt lumen forming crypt abscesses. Severe activity is indicated by neutrophils in combination with erosion or ulceration of the epithelium. Quiescent disease reflects features of chronicity, such as crypt distortion, shortening, drop-out, basal plasmacytosis, and pyloric or Paneth cell metaplasia, without any evidence of active inflammation.

**Table S3. Histologic criteria for scoring depth of chronic inflammation.**

A score of 0 indicates no chronic inflammation. A score of 1 reflects chronic inflammation confined to the mucosa. A score of 2 indicates inflammation extending into the submucosa. A score of 3 shows chronic inflammation extending from the submucosa into the muscularis propria. A score of 4 denotes transmural inflammation, with chronic inflammation extending through the muscularis propria into the subserosal or pericolonic fibroadipose tissue.

**Table S4. High-risk “non-UC” histologic features in TAC specimens differentiate IPAA failure from non-failure patients.**

The histologic features evaluated include epithelioid granulomas, ileal inflammation, knife-like ulcers, patchy disease with skip areas, right-sided disease with rectal sparing, the depth of inflammation, and the presence at least one high-risk histologic feature. *P-values* listed from Fisher’s exact test

**Table S5. CosMx quality metrics and sample information.**

Number of cells, number of FOV, mean negative control probe counts/probe/cell, and mean false code fraction/cell (see Methods), and patient information for each CosMx case.
