## Supplementary material for "Histopathologic Evaluation and Single-Cell Spatial Transcriptomics of the Colon Reveal Cellular and Molecular Abnormalities Linked to J-Pouch Failure in Patients with Inflammatory Bowel Disease": Table S1

| Table S1. Demographic and Clinical Characteristics of Case-Control Cohort | | | |
| --- | --- | --- | --- |
|  | **IPAA Non-Failure (n=39)** | **IPAA Failure (n=39)** | ***P* value** |
| Male | 20 (51%) | 19 (49%) | - |
| Female | 19 (49%) | 20 (51%) | - |
| Mean age, years (range) | 37 (13-50) | 38 (17-66) | - |
| Mean disease duration, months (range) | 98 (6-327) | 109 (3-552) | 0.68 |
| Preoperative biologics | 16 | 16 | 1.0 |
| Preoperative *C. difficile* infection | 5 | 6 | 1.0 |
| Mean follow-up time post-IPAA, months (range) | 95 (24-218) | 106 (8-198) | 0.34 |
| Mean pouch duration, months (range) | - | 55.5 (1-158) | - |
| **Indication for TAC** |  |  |  |
| Refractory to medical management | 29 (74.5%) | 24 (61%) | 0.33 |
| Fulminant colitis | 4 (10%) | 12 (31%)* | **0.048** |
| Dysplasia | 6 (15.5%) | 3 (8%) | 0.48 |
| **Stages of Operation** |  |  |  |
| 1 Stage | 9 (23%) | 9 (23%) | - |
| 2 Stages | 17 (44%) | 17 (44%) | - |
| 3 Stages | 13 (33%) | 13 (33%) | - |
| **Preoperative Diagnosis (Biopsy)** |  |  |  |
| Ulcerative colitis | 26 (67%) | 34 (87%) | 0.058 |
| Indeterminate colitis | 13 (33%) | 5 (13%) | 0.058 |
| **TAC Pathologic Diagnosis** |  |  |  |
| Ulcerative colitis | 39 (100%)* | 31 (80%) | **0.0052** |
| Biopsy-indeterminate colitis | 13 | 4 |  |
| Indeterminate colitis | 0 | 6 (15%)* | **0.025** |
| Biopsy-ulcerative colitis | - | 5 |  |
| Crohn’s disease | 0 | 2 (5%) | 0.49 |
| Biopsy-ulcerative colitis | - | 2 |  |

IPAA, ileal pouch-anal anastomosis; TAC, total abdominal colectomy
