## Supplementary material for "Histopathologic Evaluation and Single-Cell Spatial Transcriptomics of the Colon Reveal Cellular and Molecular Abnormalities Linked to J-Pouch Failure in Patients with Inflammatory Bowel Disease": Table S2

| Table S2. Histologic Criteria for Grading of Disease Activity | |
| --- | --- |
| **Activity Grading** | **Histologic Criteria** |
| Mild | Neutrophils present in epithelium |
| Moderate | Neutrophils present in crypt lumen forming crypt abscess |
| Severe | Neutrophils + erosion or ulceration of epithelium |
| Quiescent | Features of chronicity (crypt distortion/shortening/drop-out, basal plasmacytosis, pyloric or Paneth cell metaplasia) in the absence of mild/moderate/severe activity |
