## Supplementary material for "Histopathologic Evaluation and Single-Cell Spatial Transcriptomics of the Colon Reveal Cellular and Molecular Abnormalities Linked to J-Pouch Failure in Patients with Inflammatory Bowel Disease": Table S3

| Table S3. Histologic Criteria for Scoring Depth of Chronic Inflammation | |
| --- | --- |
| **Depth of Inflammation Score** | **Histologic Criteria** |
| 0 | No increase in chronic inflammation (normal) |
| 1 | Increased chronic inflammation within mucosa only |
| 2 | Increased chronic inflammation within mucosa and extension into submucosa |
| 3 | Chronic inflammation extending from submucosa into muscularis propria |
| 4 | Chronic inflammation extending from submucosa through muscularis propria and into subserosal/pericolonic fibroadipose tissue (transmural inflammation) |
