## Supplementary material for "Histopathologic Evaluation and Single-Cell Spatial Transcriptomics of the Colon Reveal Cellular and Molecular Abnormalities Linked to J-Pouch Failure in Patients with Inflammatory Bowel Disease": Table S4

| **Table S4. High-risk “non-UC” histologic features in total abdominal colectomy specimens differentiate IPAA failure from non-failure patients.**  The histologic features evaluated include epithelioid granulomas, ileal inflammation, knife-like ulcers, patchy disease with skip areas, right-sided disease with rectal sparing, the depth of inflammation, and the presence at least one high-risk histologic feature. | | | | |
| --- | --- | --- | --- | --- |
| **High-Risk Histologic Features** | **IPAA Non-Failure (n= 39)** | **IPAA Failure (n=39)** | ***P* value** | **Outcome CD**  **(IPAA Failure)** |
| Epithelioid granulomas | 0 | 1 (3%) | 1.00 | 1/1 (100%) |
| Ileal inflammation | 0 | 3 (8%) | 0.24 | 3/3 (100%) |
| Knife-like ulcers | 1 (3%) | 15 (38%)* | **0.0001** | 10/15 (67%) |
| Patchy disease with skip areas | 0 | 2 (5%) | 0.49 | 2/2 (100%) |
| Right-sided disease with rectal sparing | 0 | 1 (3%) | 1.00 | 0 |
| Inflammation depth score of 3 or more (in at least one sampled section) | 7 (18%) | 23 (59%)* | **0.0004** | 13/23 (57%) |
| At least one high-risk histologic feature present | 8 (21%) | 27 (69%)* | **<0.0001** |  |

UC, ulcerative colitis; IPAA, ileal pouch-anal anastomosis; CD, Crohn’s disease
